# Establishment of a humanized patient-derived xenograft mouse model of high-grade serous ovarian cancer for preclinical evaluation of combination immunotherapy

**DOI:** 10.1101/2025.08.08.669244

**Authors:** Luka Tandaric, Line Bjørge, Martine Rott Lode, Cecilie Fredvik Torkildsen, Pia Aehnlich, Rammah Elnour, Daniela Elena Costea, Lars Andreas Akslen, Liv Cecilie Vestrheim Thomsen, Emmet McCormack, Katrin Kleinmanns

## Abstract

The limited efficacy of immunotherapy in clinical trials in high-grade serous ovarian cancer (HGSOC) may improve by implementing models more reflective of human biology into preclinical studies. To address this, we developed and validated a humanized patient-derived xenograft mouse model of HGSOC. Human hematopoietic stem cells and patient-derived HGSOC were engrafted into immunodeficient mice. The mice were administered durvalumab and/or oleclumab intraperitoneally semi-weekly for five weeks. The immunotherapy was well-tolerated, though no responses occurred. Leukocytes in primary tumors were analyzed immunohistochemically, and circulating T cells were characterized using spectral flow cytometry. All tumors exhibited an immune-excluded immunophenotype. No significant inter-group differences in disease burden, intratumoral leukocyte density, or circulating T-cells were observed. In the durvalumab-only group, tumor burden significantly positively correlated with intratumoral cytotoxic and regulatory T-cell densities. This model reflects human disease biology and clinical findings, providing a robust platform for studying tumor-immune interactions and immunosuppressive mechanisms in HGSOC.

## 1 Introduction

High-grade serous ovarian cancer (HGSOC) is the most common and lethal subtype of ovarian cancer, with a 5-year survival rate of less than 50% [1]. The implementations of poly(ADP-ribose) polymerase inhibitors, bevacizumab and mirvetuximab soravtansine-gynx as additions to the standard treatment approach have notably improved outcomes of primary and recurrent disease [2,3]. Despite these advances, HGSOC outcomes are still frequently impaired by treatment resistance and disease recurrence [4,5], underscoring the urgent need for the exploration of more effective therapeutic approaches.

HGSOC is considered an immunogenic tumor as the vast majority of patients exhibit tumor-reactive leukocytes [6], and tumor-infiltrating leukocytes (TILs), including CD8^+^ T cells and regulatory T cells (Tregs), having a clear prognostic and therapeutic impact [7,8]. These features support the potential of immunotherapy as a viable treatment strategy for HGSOC. Although immunotherapy in the form of immune checkpoint inhibitors (ICIs) has proven effective in several solid cancer types, leading to regulatory approvals [9,10], clinical trials in HGSOC have consistently yielded disappointing results, with single-agent ICIs achieving response rates of only 10-15% [11,12]. One of the key factors responsible for immunotherapy failure in HGSOC is the immunosuppressive network within its tumor microenvironment (TME), which employs multiple immunoinhibitory mechanisms and exhibits remarkable plasticity in response to immunotherapeutic interventions, further complicating treatment efforts [13,14]. Thus, there is a need for immunotherapy combinations that can surpass these treatment resistance mechanisms of HGSOC. However, no clinical trials examining such approaches have yet demonstrated significant improvements to disease outcome [11,12,15].

These trials were often based on experiments in preclinical *in vitro* or *in vivo* models that inadequately represented the TME of HGSOC and the intricate dynamics of its interaction with the human immune system [16–21]. For example, syngeneic mouse models incorporating murine cancer cell lines implanted into mouse strains with a fully functional murine immune system have commonly been used to evaluate the effectiveness of immunotherapy in HGSOC due to their simplicity and rapid establishment [22,23], leading to a lack of translational success in clinical applications. To ensure that only biologically relevant immunotherapy combinations advance to human clinical trials, it is imperative to develop preclinical animal models that more accurately reflect the complexity of the disease.

An alternative approach to the above models is to use patient-derived xenograft (PDX) models, established by implanting tumor material from patients into immunodeficient mice. This type of model effectively preserves the structural and genomic features of the primary tumor, as well as intratumoral heterogeneity, and treatment response [24,25]. While PDX models are typically established via subcutaneous or intraperitoneal injection, orthotopic PDX models, in which patient tumor material is engrafted into the corresponding anatomical site in the model animal, offer the most biologically representative approach as they also preserve the pattern of metastatic spread [26]. However, orthotopic PDX models are less frequently utilized due to their highly immunodeficient nature, requiring specialized handling facilities and procedures, as well as the need for advanced surgical expertise and the reliance on imaging techniques for tumor growth monitoring [27].

Despite facilitating improved representation of the HGSOC TME in the preclinical setting, to effectively simulate the tumor-immune cell interactions critical for disease outcome and immunotherapy success [7,8,28], orthotopic PDX models require the co-engraftment of a human immune system. Humanization of mice can be accomplished through the adoptive transfer of allogenic or autologous leukocyte subsets, most commonly peripheral blood mononuclear cells or T cells. However, these approaches provide only partial humanization, as they fail to capture the full complexity of the human immune system. Furthermore, these leukocytes, having undergone maturation, are more likely to induce graft-versus-host disease when placed in a non-human environment, such as a mouse [29–31]. In contrast, injection of CD34^+^ human hematopoietic stem cells (HSCs) generates a more comprehensive human immune system, eliminating the risk of graft-versus-host disease by resulting in the development of a variety of human leukocytes adapted to murine physiology [32,33]. Thorough humanization is most consistently established in heavily immunodeficient strains, such as the *NOD.Cg-Prkdc^scid^ Il2rg^tm1Wj/^SzJ* (NSG) and *NOD.Cg-Prkdc^scid^ Il2rg^tm1Wjl^ Tg (CMV-IL3, CSF2, KITLG) 1Eav/MloySzJ* (NSGS) strains [34,35]. Our group has previously successfully developed HSC-humanized orthotopic PDX mouse models of HSGOC which have been used to accurately replicate the low efficacy of single-agent nivolumab in HGSOC [33].

The NSGO-OV-UMB1/ENGOT-OV30 clinical trial, which evaluated combined durvalumab (anti-PD-L1) and oleclumab (anti-CD73) immunotherapy in HGSOC [15], reported a response rate of 4%, which is consistent with the generally low response rates to immunotherapy observed in HGSOC [11,12]. One motivation for testing this combination in a clinical trial on HGSOC was a successful preclinical study involving treatment of a non-humanized BALB/c mouse model of murine colorectal cancer with a combination of oleclumab and a murine-PD-1-binding antibody that, analogous to the mechanism of durvalumab, blocks PD-1/PD-L1 interaction. However, in contrast to the results of the NSGO-OV-UMB1/ENGOT-OV30 clinical trial, this preclinical study reported widespread tumor rejection in model mice [21]. To our knowledge, the durvalumab-oleclumab combination has not been validated in preclinical model systems that more accurately replicate the HGSOC TME or its interactions with the human immune system. The aim of this study was to advance preclinical testing of immunotherapy combinations in HGSOC by developing a biologically relevant immunocompetent mouse model of the disease and validating its fidelity by emulating the NSGO-OV-UMB1/ENGOT-OV30 clinical trial.

Here, we present our humanized orthotopic PDX mouse model of HGSOC, which accurately replicates the morphology, immune contexture, and immunotherapy resistance of the immunosuppressive HGSOC TME. Using this model, we have successfully replicated the conditions and outcome of the NSGO-OV-UMB1/ENGOT-OV30 clinical trial. Our results address the critical need for representative preclinical models for testing combination immunotherapy in HGSOC by providing a robust preclinical platform that can enhance the reliability of preclinical data and contribute to the improvement of the design and outcomes of future clinical trials.

## 2 Materials and Methods

### 2.1 Ethical considerations concerning experimental model animals

This study was conducted in compliance with the procedures outlined by the Norwegian State Commission for Laboratory Animals and with the approval of the Norwegian Food Safety Authority (Application ID: 25412). Female NSGS mice (aged 6–12 weeks) (Cat.No. 013062, The Jackson Laboratory, USA), bred at the animal facility of the University of Bergen, were housed in groups of up to five mice in individually ventilated HEPA-filtered cages, with regular replacement of autoclaved food, water, bedding and cages.

### 2.2 Acquisition and processing of human tumor material

For this study, an International Federation of Gynecology and Obstetrics (FIGO) stage IIa HGSOC tumor from a treatment-naïve patient was provided by the Gynecologic Cancer Biobank, Women’s Clinic, Haukeland University Hospital, Bergen, Norway. Ethical approval (REK ID: 2014/1907, 2017/612) and written informed consent from the patient were obtained prior to tumor tissue collection. The tumor had wild-type *BRCA1/2* and was later classified as platinum-resistant. The tumor was sampled during primary cytoreductive surgery and dissociated as previously described [33]. Single cells (henceforth referred to as “PDX material”) were cryopreserved at −150°C in a mix of 90% V/V fetal bovine serum (FBS) (Cat.No. F7524, Sigma-Aldrich, USA) and 10% V/V dimethyl-sulfoxide (Cat.No. D8418, Sigma-Aldrich, USA). This PDX material was selected because it exhibited the highest proportion of CD73-expressing tumor cells among the HGSOC PDX models in our model portfolio (Fig. S1) [36], aligning with the inclusion criterion of the NSGO-OV-UMB1/ENGOT-OV30 clinical trial according to which over 10% of tumor cells need to be CD73-positive [15]. The PDX material was propagated by two rounds of *in vivo* passaging in immunodeficient mice. Thawed PDX material was implanted into the ovarian bursae of the mice (described in detail in section 2.6), and the resulting tumors were harvested, processed and cryopreserved in the same manner as the original patient material.

### 2.3 Lentiviral transduction of tumor cells

To enable the *in vivo* monitoring of tumor growth and metastasis in model mice using bioluminescence imaging (BLI), PDX material was transduced using RediFect Red-FLuc-GFP lentiviral particles (Cat.No. CLS960003, PerkinElmer, USA), containing genes encoding *Luciola italica* firefly luciferase (luc) and green fluorescent protein (GFP) reporters. The transduction was performed according to the following custom protocol. Twice-passaged PDX material was thawed and washed by centrifugation (400 g, 5 min, room temperature (RT)) in RPMI 1640 cell culture medium (Cat.No. R5886, Sigma-Aldrich, USA) supplemented with 10% V/V HyClone FBS (Cat.No. SH30071.03HI, Cytiva, USA), 1% V/V L-glutamine (Cat. No. G7513, Sigma-Aldrich, USA) and 1% V/V penicillin-streptomycin (Cat.No. P0781, Sigma-Aldrich, USA) (henceforth referred to as “complete RPMI medium”). Cells were counted, seeded into an adherent cell culture plate (Cat.No 3538, Corning, USA) at a pre-determined number per well, and incubated in complete RPMI medium (18-24h, 37°C, 5% V/V CO_2_). After incubation, to establish a precise multiplicity of infection (MOI) of 20, cells in pre-specified wells were re-counted - the medium was aspirated, and the cells were detached from the plate after a 5-20 minute incubation in 100 µL of accutase (Cat.No. SCR005, Sigma-Aldrich, USA), neutralized by the addition 1 mL of complete RPMI medium. The medium in the remaining wells was aspirated and replaced with complete RPMI medium containing no FBS and supplemented with Vectofusin-1 (Cat.No. 130-111-163, Miltenyi Biotec, Germany) to enhance transduction efficiency. Lentiviral particles were added to the wells at an MOI of 20, followed by spinoculation, which involved centrifugation of the cell culture plate (700 g, 90 min, 32°C) and subsequent incubation (16-24 h, 37°C, 5% V/V CO_2_). After incubation, the plate was centrifuged (400 g, 5 min, RT), and the culture medium containing lentiviral particles was aspirated and replaced with phosphate-buffered saline (PBS). After centrifugation of the plate (400 g, 5 min, RT) and aspiration of the PBS, cells were detached using accutase as described earlier, transferred to 1.5 mL tubes, and centrifuged (400 g, 5 min, RT). Each cell pellet was subsequently washed by resuspension in PBS and centrifugation (400 g, 5 min, RT). Cells were then orthotopically implanted into mice (described in detail in section 2.6). Transduced PDX material was propagated *in vivo* for three passages. During each passage, excised tumors were dissociated and cryopreserved as described beforehand, and subsequently thawed PDX material was enriched for high GFP expression prior to the next orthotopic injection using a high-speed cell sorter (Model SH800, Sony Biotechnology, USA).

### 2.4 Isolation of human hematopoietic stem cells

The CD34^+^ human HSCs used in this study were isolated from umbilical cord blood collected during cesarean delivery of a healthy woman with a presumedly healthy pregnancy (Research Biobank for Blood Diseases, Haukeland University Hospital, Bergen, Norway). Ethical approval (REK ID: 2015/1759) and written informed consent from the parents were obtained prior to blood collection. The umbilical cord was clamped, and the distal portion of the umbilical vein was immediately punctured with a 12-gauge needle. A volume of 140 mL of blood was collected in a collection bag containing citrate phosphate dextrose anticoagulant solution (Cat.No. MSC1208DU, Macopharma, France). Mononuclear cells (MNCs) were then isolated from the umbilical cord blood as follows: The blood was diluted 1:1 with sterile, room-temperature PBS. Diluted blood was laid on top of a density gradient medium - Lymphoprep (Cat.No. #07861, STEMCELL Technologies, Canada), and centrifuged (400 g, 30 min, RT, lowest acceleration, no brakes). Opaque interphases containing MNCs were collected into sterile conical 50 mL tubes and washed by resuspension in a large volume of PBS and subsequent centrifugation (500 g, 5 min, RT). The supernatant was aspirated and the remaining red blood cells (RBCs) were lysed by incubating the cells in 20 mL of 1x RBC Lysis Buffer (Cat.No. TNB-4300, Cytek Biosciences, USA) (8 min, RT, shielded from light). The cell suspension was supplemented with magnetic activated cell sorting (MACS) buffer (2 mM EDTA and 0.5% w/V bovine serum albumin in PBS, pH 7.2) to a total volume of 45 mL and centrifuged (300 g, 10 min, RT). After aspiration of the supernatant, the MNCs were resuspended in 1 mL of CryoStor CS10 (Cat.No. 210502, Biolife Solutions, USA) and cryopreserved at −150°C. On the day of mouse humanization, the MNCs were thawed at 37°C. All MNCs were combined in a new sterile conical 50 mL tube and CD34^+^ HSCs were isolated using MACS with the CD34 MicroBead Kit (Cat.No. 130-046-702, Miltenyi Biotec, Germany), according to the manufacturer’s protocol. The purity of isolated CD34^+^ HSCs was determined by staining an aliquot of the CD34-enriched cell suspension with a phycoerythrin-(PE) labeled anti-human-CD34 antibody (Cat.No. 130-098-140, Miltenyi Biotec, Germany) and analyzing the cells with an LSRFortessa Cell Analyzer (Cat.No. 649225, Becton, Dickinson and Company, USA) equipped with BD FACSDiva Software (v9.0.1, Becton, Dickinson and Company, USA) (Fig. S2). Detailed protocols for HSC isolation and purity control are available as supplementary files.

### 2.5 Humanization of model mice

The NSGS mice were humanized by injection of 1.9 × 10^4^ CD34^+^ HSCs suspended in 100 µL of saline into the tail vein (Fig. 1A). All mice were injected with HSCs from the same donor. Chimerism was evaluated by conventional flow cytometry as previously described [33] at three timepoints following HSC injection: eight weeks (before orthotopic PDX implantation), 11 weeks (prior to treatment initiation) and 17 weeks (at the study endpoint) (Fig. 1A; Fig. S3).

**Fig. 1.**
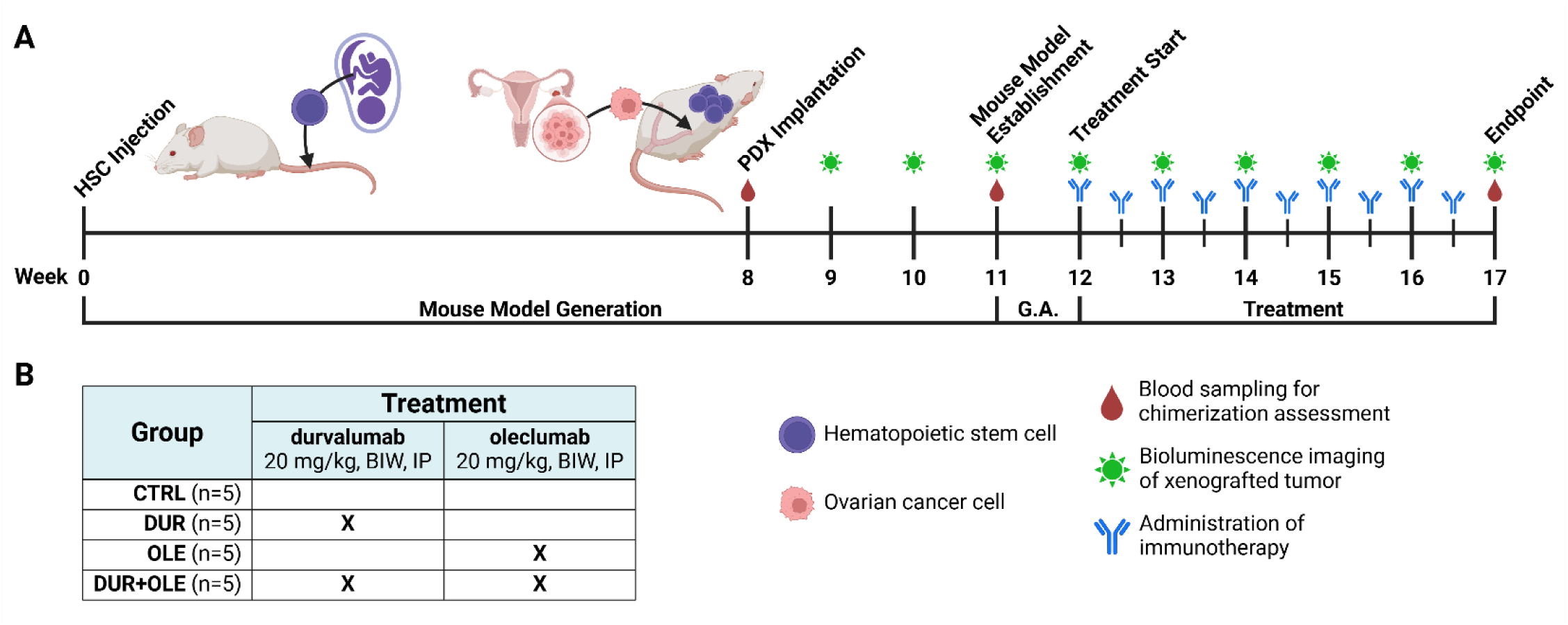
Establishment and application of the mouse model. (A) Timeline for the generation, monitoring, and combination immunotherapy treatment of a murine orthotopic PDX model of treatment-naïve high-grade serous ovarian cancer. (B) Distribution of model mice and description of immunotherapy administration across treatment groups. HSC - hematopoietic stem cell; PDX - patient-derived xenograft; G.A. - group assignment; BIW - twice a week; IP - intraperitoneally; CTRL - control group; DUR - durvalumab-only group; OLE - oleclumab-only group; DUR+OLE - combination treatment group

### 2.6 Establishment of high-grade serous ovarian cancer xenografts in humanized mouse models

To establish the orthotopic PDX mouse model, eight weeks after HSC injection, GFP^+^/luc^+^ PDX material (passage five) was thawed, and 1.0 × 10^5^ cells were orthotopically injected into the bursa of the right ovary of each of 20 NSGS mice, as described previously [37] (Fig. 1A). Briefly, GFP^+^/luc^+^ cells were prepared for orthotopic injection by resuspension in saline, followed by mixing two parts of the cell suspension with one part of a 1:1 mix of Matrigel membrane matrix (Cat.No 10365602, Corning, USA) and RPMI1640 cell culture medium. After the mice were administered analgesia (5 mg/kg meloxicam (Metacam, 2 mg/mL injection solution, Cat.No. 386860, Boehringer Ingelheim, Germany) and 0.1 mg/kg buprenorphine hydrochloride (Temgesic, 0.3 mg/mL injection solution, Cat.No. 521634, Indivior Inc., USA), anesthesia and ophthalmic lubricant, a small incision (∼5 mm) was made through the skin and abdominal muscles, mid-way between the last rib and the iliac crest. The right ovary was grasped by its surrounding fat pad and exteriorized through the incision. With the aid of a microscope with a 10x magnification (SMZ-171, Motic, China), a 30-gauge needle syringe was used to inject 10 µL of the PDX material suspension into the ovarian bursa. Matrigel was allowed to polymerize by delaying the removal of the needle, preventing leakage of the cell suspension. The ovary was carefully repositioned, and the incision was sutured using an absorbable suture (Cat.No. J492G, Agntho’s, Sweden). Postoperatively, the animals were given a subcutaneous injection of sterile saline and were allowed to recover in a warm environment before being returned to their home cage.

Growth of the luc^+^ PDX material was monitored weekly by BLI (Fig. 1A). Mice were injected intraperitoneally with 150 mg/kg of D-luciferin (Cat.No. L-8220, Biosynth, Switzerland). Bioluminescent signal was acquired laterally and ventrally 10 minutes after D-luciferin administration using the IVIS Spectrum In Vivo Imaging System (Perkin Elmer, USA). Images were analyzed using Living Image software (v4.7.3, Perkin Elmer, USA).

### 2.7 Treatment

Humanized PDX mice were stratified into four treatment groups containing mice with similar distributions of leukocyte chimerism extent and tumor load, as determined by conventional flow cytometry and BLI, respectively. Based on the NSGO-OV-UMB1/ENGOT-OV30 clinical trial of durvalumab and oleclumab in HGSOC [15], and the pre-clinical study by Hay et al. [21], the groups and drug dosages were defined as follows: (a) 20 mg/kg durvalumab (n=5); (b) 20 mg/kg oleclumab (n=5); (c) 20 mg/kg of both durvalumab and oleclumab (n=5); (d) untreated control (n=5) (Fig. 1B). Durvalumab (EU No. EU/1/18/1322/002, batch AAUR, AstraZeneca, UK) and oleclumab (Cat.No. HY-P99039, batches 279777 & 255720, MedChemExpress, USA) were diluted in sterile saline to concentrations of 3.8 mg/mL and 5.0 mg/mL, respectively, and administered intraperitoneally twice per week for five subsequent weeks. The appearance, activity levels, and food and water intake of the mice were monitored daily, and their weight was measured multiple times a week. Tumor growth was monitored by BLI. Humane endpoints were defined using score sheets and were based on weight loss of over 10% since the most recent weighing, distension of the abdomen caused by ascites, unkempt fur, paleness or lethargy. During the first week of treatment, two mice from the oleclumab-only group died. In order to preserve statistical power, one mouse from the control group was re-assigned to the oleclumab-only group and was administered oleclumab from the second treatment timepoint onwards.

### 2.8 Sample collection and processing

At the end of the study, mice were euthanized according to institutional guidelines due to a combination of factors, including health deterioration. Briefly, mice were anesthetized using sevoflurane (Cat.No. 002185, Vitusapotek, Norway). While under anesthesia, a terminal blood sample was collected from the facial vein into an EDTA Microvette (Cat.No. 20.1341.100, Sarstedt, Germany), followed by cervical dislocation. The collected blood was processed using Stable-Lyse2 (Cat.No STBLYSE2-250, Smart Tube, USA) and Stable-Store2 (Cat.No STBLSTORE2-1000, Smart Tube, USA), according to the manufacturer’s protocol, and then frozen in cryogenic vials at −80°C. Euthanized mice were dissected ventrally, and the primary tumor characteristics, as well as the presence and extent of metastases, were described macroscopically. Samples of the primary tumors were fixed in 4% V/V formaldehyde (Cat.No. 9713.9010, VWR International, USA) for 24 hours, then washed using deionized water and kept in 70% V/V ethanol until paraffinized and sectioned for immunohistochemical (IHC) analysis.

### 2.9 Spectral flow cytometry analysis of the peripheral blood of model mice

Lysed and fixed samples of whole blood were thawed, washed, and stained for spectral fluorescence flow cytometry analysis. A detailed protocol, as well as the antibody panel (Table S1), are available as supplementary files. Briefly, blood samples were thawed at 4°C, mixed with 0.25 mg/mL DNAse I (Cat.No. DN25, Sigma-Aldrich, USA) in Dulbecco’s PBS containing Ca^2+^ and Mg^2+^ (Cat.No. D8662, Sigma-Aldrich, USA), and supplemented with CountBright Absolute Counting Beads (Cat.No. C36950, ThermoFisher Scientific, USA). Samples were washed in PBS and filtered through the 40 µm mesh in the cap of 5 mL round-bottom tubes (Cat.No. 352235, Corning, USA). Pelleted cells were incubated in a solution of human Fc receptor blocking agent (Cat.No. 130-059-901, Miltenyi Biotec, Germany) and anti-mouse-CD16/CD32 monoclonal antibody (Cat.No. 16-0161-82, ThermoFisher Scientific, USA), and then stained with the antibody mix defined in Table S1. Stained cells were washed twice with a mix of 2% V/V FBS in PBS and acquired on an ID7000 Spectral Cell Analyzer (LE-ID7000C, Sony Biotechnology, USA) equipped with ID7000 Software (v2.0.2.17121, Sony Biotechnology, USA). Single-cell data was processed using FlowJo software (v10.10.0, Becton, Dickinson and Company, USA) (Fig. S4). Absolute leukocyte quantities were calculated using blood volume estimations based on Counting Bead data. The composition of the human leukocyte pool in each sample was calculated by dividing the number of cells of a specific subset by the total number of human leukocytes in that sample.

### 2.10 Immunohistochemical staining of primary mouse tumors

Immunohistochemical staining of serial sections of paraffinized primary tumor tissue was performed according to the protocol available as a supplementary file. In short, paraffinized tumor sections with a thickness of 3 µm were deparaffinized in xylene and rehydrated in a graded ethanol series. After antigen retrieval at a pH of 9.0, blocking of endogenous peroxidase activity was performed. Next, tissues were incubated for 45-60 minutes in a blocking solution of 3% w/V bovine serum albumin (Cat.No. 10735086001, Merck, USA) to mitigate non-specific antibody binding. After two washes, primary antibodies targeting human CD45 (Cat.No. 14-9457-82, ThermoFisher Scientific, USA), CD20 (Cat.No. 555677, Becton, Dickinson and Company, USA), CD3 (Cat.No. ab17143, Abcam, UK), CD8 (Cat.No. 372902, BioLegend, USA) and FoxP3 (Cat.No. 14-4777-82, ThermoFisher Scientific, USA) were individually applied to five serial sections of each primary PDX tumor and left to incubate overnight at 4°C. Antibody details are provided in Table S2. The next day, primary antibodies were washed off, and appropriate horseradish-peroxidase-labeled secondary antibodies were applied to the tissues. After the secondary antibody was washed off, antigen localization was visualized via a 3,3’-diaminobenzidine reaction. Stained tissues were washed, counterstained with hematoxylin (Cat.No S3301, Agilent, USA), and mounted using an automated coverslipper (Model 4740, Sakura, Japan). The immunophenotype [38] of all primary tumors was microscopically evaluated as immune excluded by a trained pathologist, as leukocytes were present within the tumors, but confined to the stroma surrounding tumor cell foci.

### 2.11 Digital tissue analysis

Stained primary tumor slides were digitally scanned at a 400x magnification using a slide scanner (Model BX61VSF, Olympus, Japan) equipped with Olympus VS-ASW software (v2.9.2, Olympus, Japan). Digital cell segmentation and quantification were performed on the high-resolution digital scans using QuPath software (v0.5.1) [39]. Briefly, based on previously published approaches [40,41], areas with the highest density of leukocyte infiltration were selected for annotation. Due to the immune-excluded nature of the tumors, annotations were drawn along the invasive margin (IM) of tumors, symmetrically extending 200 µm from each side of the tumor-stroma border of the IM. Automatic staining vector estimation and optical-density-based cell segmentation were used for the enumeration of marker-positive leukocytes. A detailed overview of positive cell detection parameters is available in Table S3. For each tumor and marker, the density of marker-positive TILs per mm^2^ was calculated by dividing the total number of positive TILs by the total combined size of the annotated area.

### 2.12 Statistical analyses

Statistical analyses were performed using GraphPad Prism (v10.4.1, GraphPad Software, USA). Primary tumor volumes at endpoint, quantities of leukocytes per volume of blood, relative leukocyte abundances, and TIL densities were compared between treatment groups. For each sample group (n=4 or n=5), the Shapiro-Wilk test was used to assess the normality of the datasets. Inter-group comparisons were performed using the Kruskal-Wallis test with Dunn’s multiple comparison correction. Ratios of CD8^+^ and FoxP3^+^ TIL densities were compared between groups using one-way ANOVA with Tukey’s multiple comparison correction. Correlations between TIL density and tumor burden (volume) were evaluated as follows: for each staining marker, measurements of leukocyte density were either stratified according to treatment group or considered as one group. After assessment of dataset normality using the Shapiro-Wilk test, Pearson or Spearman rank correlation tests were performed. Correlation plots were modeled using simple linear regression. Statistical significance was defined as p < 0.05.

## 3 Results

### 3.1 Successful establishment of a humanized PDX mouse model for pre-clinical testing of combination immunotherapy in HGSOC

We successfully implemented the mouse model establishment workflow described by Kleinmanns et al. [33], utilizing HSCs and tumor material from unique donors to create a humanized PDX model for pre-clinical testing of combination immunotherapy in HGSOC (Fig. 1A). Despite injecting significantly fewer CD34^+^ HSCs per mouse compared to Kleinmanns et al., chimerism analysis of peripheral blood samples obtained at multiple timepoints after the injection demonstrated stable HSC engraftment and sustained development of human lymphocyte populations in the experimental mice (Table S4). Following orthotopic implantation of PDX material in week eight, BLI conducted during weeks 9-11 revealed detectable tumor signal localized to the ovary in all mice, confirming successful orthotopic tumor engraftment (Table S5). Following confirmation of sufficient chimerism and stable PDX engraftment, the 20 experimental mice were evenly allocated into four treatment groups, ensuring comparable mean chimerism levels and bioluminescence signal intensities across groups (Fig. 1B). All mice receiving combination therapy demonstrated good tolerance to the treatment, with no observable deterioration in their condition compared to the other treatment and control groups.

### 3.2 The established HGSOC PDX tumor model resists growth inhibition by immunotherapy

Longitudinal weekly BLI of the xenografted PDX material showed consistent tumor growth and progressive disease in all mice (Fig. 2A, Fig. S5). Although the average bioluminescent signal was marginally higher in the control group than in all treatment groups throughout the observation period (Fig. 2B), this difference was not statistically significant. Furthermore, tumor growth kinetics were similar across all four groups, with none of the treatments resulting in a sustained reduction in bioluminescence signal (Fig. 2B). At the end-of-study necropsy, measurements of excised primary tumor volume showed no statistically significant difference in tumor burden between the treatment groups (Fig. 2C, Table S6). The extent of metastatic dissemination was also similar across groups, with nearly all mice exhibiting abdominal carcinomatosis, with visible metastatic lesions on the omentum, peritoneal wall, and/or diaphragm (Table S7).

**Fig. 2.**
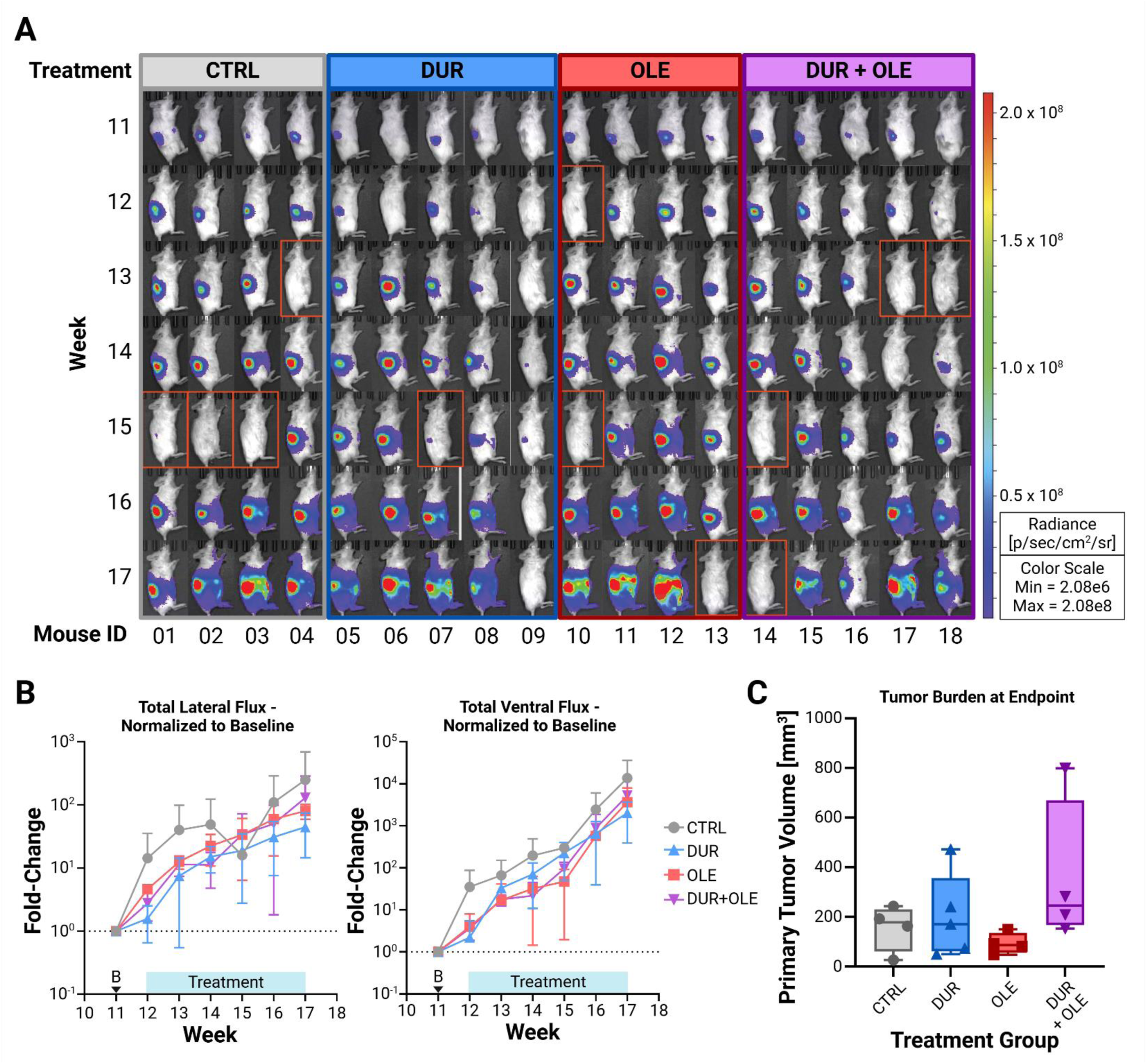
Monitoring of tumor burden in PDX-implanted experimental mice. (A) Longitudinal overview of weekly bioluminescence imaging results for PDX-implanted mice in the lateral position. Each column represents an individual mouse. Only images from baseline (week 11) through endpoint (week 17) are shown. Mice outlined with an orange border displayed inexplicably low bioluminescence at the specified timepoint, even after luciferin re-injection. These data were excluded from further analyses. Bioluminescence images of the mice taken ventrally are displayed in Fig. S5. Full data on the total lateral flux are available in Table S5. (B) Average total lateral (left graph) and ventral (right graph) photon flux in each treatment group during treatment, relative to baseline (marked by a “B” on the X-axis). (C) Comparison of tumor burden at the end of the study between treatment groups. Only the primary tumor was included in tumor burden assessment due to the small size of the metastatic lesions. Tumor volume data is available in Table S6. CTRL - control group; DUR - durvalumab-only group; OLE - oleclumab-only group; DUR+OLE - combination treatment group

### 3.3 The administration of immunotherapy does not enhance intratumoral infiltration of immune cells in the established HGSOC PDX model

To assess the effectiveness of the combined durvalumab-oleclumab treatment in promoting leukocyte infiltration into the HGSOC PDX tumor and compare it with the impact of the individual immunotherapeutic agents, all excised primary tumors were processed for paraffin embedding, sectioned and immunohistochemically stained. The staining targeted markers for total human leukocytes (hCD45), B cells (CD20), and the intended targets of durvalumab and oleclumab immunotherapy: total T cells (CD3), cytotoxic T cells (CD8), and Tregs (FoxP3). Brightfield microscopy scans of the stained PDX tumor sections showed that, in all tumors, marker-positive cells were predominantly localized to the outer, stromal region of the IM, with scarce leukocyte infiltration into tumor-rich areas (Fig. S6). Consequently, all tumors were categorized as immune-excluded [38]. The most densely infiltrated regions of the IMs were selected for enumeration of human leukocytes using automated detection of marker-positive cells (Fig. 3A). The analysis showed no significant differences in intratumoral leukocyte infiltration among the treatment groups (Fig. 3B, Table S8). Similarly, the median ratios of cytotoxic to regulatory T cells showed no significant variation between the groups (Fig. 3C). To evaluate the relationship between leukocyte infiltration and tumor burden in the context of each treatment, we examined correlations between primary tumor volume at the end of the study and the densities of marker-positive leukocytes in the IM. No significant correlations or trends were observed in the combination treatment group or for most markers in the other groups. The only exception was the durvalumab monotherapy group, in which increased densities of CD8^+^ cells and FoxP3^+^ cells were significantly positively correlated with larger tumor burden (Fig. 3D, Fig. S7, Table S9).

**Fig. 3.**
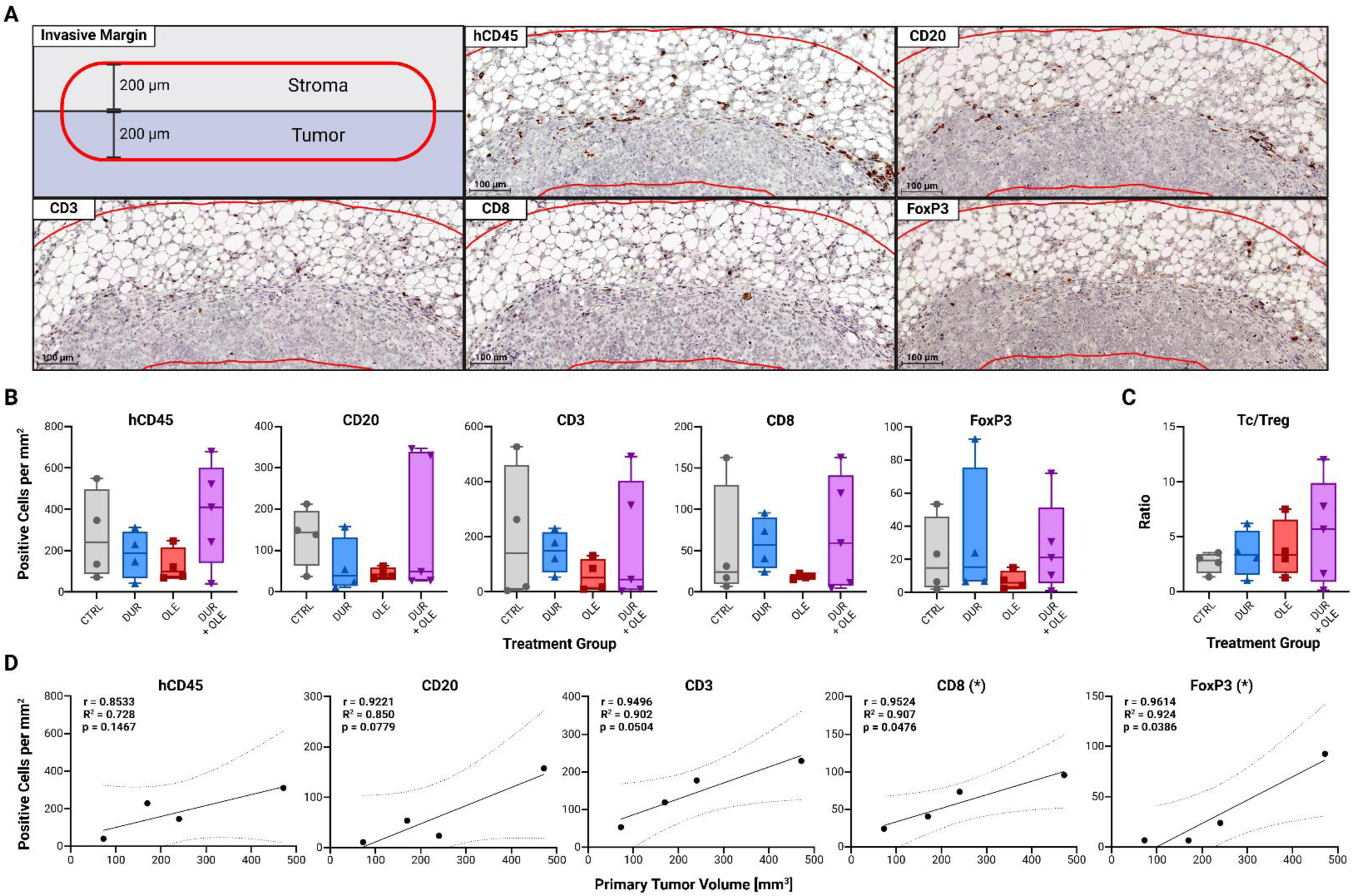
Intratumoral leukocyte densities determined by immunohistochemistry (IHC), and digital analysis using QuPath software. (A) Representative images of tumor areas rich in leukocytes selected for positive cell quantification. The top-left image is a schematic representation of the annotation method: By tracing the tumor-stroma border of the invasive margin with a 400-µm-thick brush tool, an invasive tumor margin of symmetrical intra-stromal and intra-tumoral depths of 200 µm was delineated. The remaining images display serial primary PDX tumor sections stained with antibodies targeting the leukocyte marker specified in the upper-left corner of each image. Antibody details are provided in Table S2. (B) Comparison of intratumoral leukocyte densities between treatment groups. Full data on intratumoral leukocyte density is available in Table S8. (C) Inter-group comparison of the ratios of CD8^+^/cytotoxic (Tc) and FoxP3^+^/regulatory (Treg) TIL densities. (D) Plots depicting correlations between tumor burden at the end of the study and the densities of intratumoral marker-positive leukocytes for the group of mice treated with durvalumab. Significant correlations are marked with an asterisk (*) in the graph title. All correlation plots are displayed in Fig. S7, and full correlation data is available in Table S9. CTRL - control group; DUR - durvalumab-only group; OLE - oleclumab-only group; DUR+OLE - combination treatment group; Tc - cytotoxic (CD8^+^) T cell; Treg - regulatory (FoxP3^+^) T cell; r - Pearson’s correlation coefficient

### 3.4 T-cell abundances in the peripheral blood of the humanized HGSOC PDX model mice are not altered by immunotherapy

Comprehensive characterization of the T-cell repertoire in peripheral blood collected from experimental mice at the end of the study was conducted using spectral flow cytometry. This analysis aimed to identify differences in the absolute and relative abundances of T-cell subsets across treatment groups (Fig. S4A; Table S10). The implementation of fluorescent counting beads at the start of the staining workflow enabled the precise assessment of the blood volume underlying each dataset, allowing for the determination of absolute leukocyte counts per microliter of blood. In all samples, CD4^+^ T cells exhibited higher absolute abundance than CD8^+^ T cells, with central memory cells being the most prevalent subset within both T-cell types (Fig. 4A, Table S11). While effector memory cells constituted the second most abundant CD4^+^ T-cell subset in nearly all samples, precursor effector CD4^+^ T cells were rare (Table S11). Effector and effector memory CD8^+^ T cells were extremely rare, with fewer than one cell per microliter of blood, even in samples with a high overall abundance of CD8^+^ T cells (Table S11). No inter-group differences in absolute leukocyte counts were observed (Fig. 4A). The relative abundances of T-cell subsets, normalized to total leukocytes, mirrored the patterns seen in absolute counts and showed no significant immunotherapy-induced changes (Fig. 4B; Table S12). We attempted to assess T-cell exhaustion by measuring PD-L1 expression with a previously validated antibody; however, the combination of negligible PD-L1 expression and low T-cell counts in several samples precluded reliable analysis (Fig. S4A-C; Table S13).

**Fig. 4.**
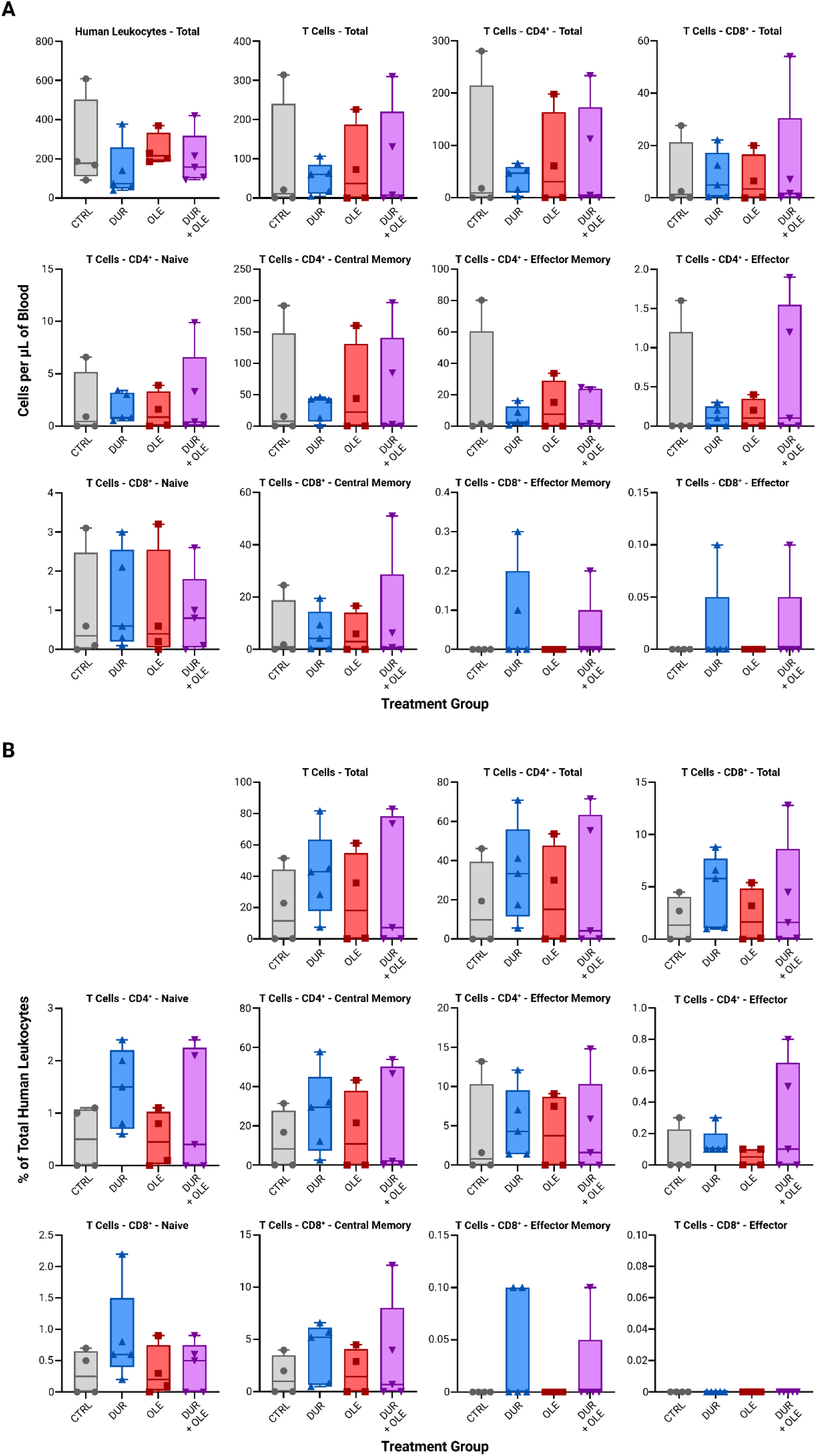
Results of the spectral flow cytometry analysis of blood samples taken at the end of the study. Data on the human leukocyte counts are available in Table S10. (A) Comparison of total human leukocyte and T-cell subset frequencies per volume of blood between treatment groups. Data on the human leukocyte counts per µL of blood are available in Table S11. (B) Distribution of T-cell subsets across treatment groups, relative to total human leukocytes. Data on the abundances of human leukocyte subsets relative to total human leukocytes in the blood are available in Table S12. CTRL - control group; DUR - durvalumab-only group; OLE - oleclumab-only group; DUR+OLE - combination treatment group

## 4 Discussion

Optimizing translation of preclinical research into clinical practice is essential for improving HGSOC outcomes. While mouse models have advanced our understanding of HGSOC biology and have become indispensable for cancer therapy development, translating preclinical efficacy into clinical outcomes remains challenging. In this study, we successfully developed a humanized orthotopic PDX mouse model and demonstrated its application in preclinical evaluation of combination immunotherapy in HGSOC by closely mirroring the setup of the NSGO-OV-UMB1/ENGOT-OV30 clinical trial.

This study builds directly on our prior development of immunocompetent mouse models, including the creation of the first HSC-humanized orthotopic PDX models of HGSOC, which enabled the characterization of tumoral and immunological responses to single-agent nivolumab [33]. Given the limited efficacy of single-agent immunotherapy in HGSOC, combination approaches based on ICIs are being tested in clinical trials. The NSGO-OV-UMB1/ENGOT-OV30 clinical trial resulted in poor response rates of HGSOC to combined durvalumab and oleclumab therapy [15]. This is in contrast to the results of the preclinical study of the drug combination by Hay et al., which had demonstrated frequent tumor rejection [21]. To investigate if the discordance between the preclinical and clinical results was due to the use of models that inadequately represented the disease *in vivo*, we sought to test the durvalumab-oleclumab regimen in our HSC-humanized orthotopic HGSOC PDX model system. The PDX tumors in our previously established immunocompetent mouse models of HGSOC did not express CD73. Since positive intratumoral CD73 expression was a key inclusion criterion for the clinical trial, we screened our patient tumor material portfolio and selected the HGSOC with the highest tumor cell CD73 expression to construct the model for this study. Determining whether the resulting tumor maintained the high expression level of CD73 throughout its establishment and growth in the experimental mice would have benefited treatment response evaluation. Unfortunately, due to technical limitations posed by small tumor size (Fig. 2A) and the ethical obligation to minimize animal suffering, a pre-treatment biopsy was not feasible.

Hay et al.’s preclinical study in a syngeneic mouse model of colorectal cancer was pivotal in initiating clinical trials for the durvalumab-oleclumab regimen in solid tumors. The study demonstrated that simultaneous inhibition of PD-1/PD-L1 interaction and adenosine generation led to tumor rejection in 60% of subcutaneous-tumor-bearing mice receiving the combined treatment [21]. Conversely, this dual-targeting strategy failed in clinical trials for ovarian, colorectal, lung and pancreatic cancer [15,42]. A possible explanation is the limited clinical translatability of Hay et al.’s preclinical study, which modelled a murine immune response to murine cancer rather than a human immune response to human malignancies. To address this, we used HSC-humanized NSGS mice orthotopically engrafted with patient-derived HGSOC tumors to evaluate durvalumab-oleclumab treatment.

The lack of discernable therapeutic benefit from all treatments in our study is in alignment with previous findings that HGSOC often exhibits resistance to immunotherapy [11,12], and with the low response rate observed in the NSGO-OV-UMB1/ENGOT-OV30 clinical trial [15]. Immunoprofiling of the primary PDX tumors showed an absence of significant differences in TIL densities among treatment groups, with all xenografts giving rise to an immune-excluded tumor immunophenotype, regardless of treatment modality. A number of preclinical studies in humanized mouse models of HGSOC, including our previous work, have reported post-treatment intratumoral T-cell infiltration following PD-1/PD-L1 interaction inhibition. However, none of them have contextualized their findings with regard to the spatial distribution of T cells within the TME, i.e., the tumor’s immunophenotype. As these studies have observed a wide variety of treatment responses despite confirming the presence of TILs, it is clear that this was a critical oversight in evaluating their models’ predictive power [31,33,43]. This, in concert with our current study’s results and previous findings that the HGSOC immunophenotype is prognostic of disease outcome and predictive of immunotherapy efficacy [44,45], emphasizes the need for evaluating the immunosuppressive capabilities of an HGSOC before its implementation into a PDX model or reaching either a preclinical or clinical decision on immunotherapy administration. This evaluation could be performed using simple IHC on tumor biopsies and would help anticipate the model’s response to immunotherapy, as well as confirm its fidelity in representing the original tumor’s characteristics. Comparing the structural phenotype and behavior of primary PDX tumors to the donor material could have further validated our model’s ability to replicate HGSOC biology.

A paradoxical observation in our study was the significant positive correlation between larger tumor burden and increased intratumoral densities of both CD8^+^ cytotoxic T cells and FoxP3^+^ Tregs in the durvalumab monotherapy group (Fig. 3D). We hypothesize that durvalumab initially enhanced CD8^+^ TIL density by facilitating their recruitment from the circulation, as well as their proliferation. The influx of CD8^+^ TILs led the FoxP3^+^ Tregs present in the TME to respond by increasing their abundance and immunosuppressive capacity, counteracting immune-mediated tumor growth suppression. This type of Treg-mediated immunosuppression during PD-1/PD-L1 interaction inhibition within an immunogenic tumor has been described in melanoma by Geels et al. Similarly to our study, they observed intratumoral accumulation of both CD8^+^ T cells and Tregs upon administration of a PD-1-targeting antibody. They demonstrated that the secretion of interleukin-2 by non-exhausted CD8^+^ T cells induces inducible costimulator (ICOS) expression in Tregs, whose ligation by ICOS-ligand in the TME stimulates activation, proliferation, and expression of T-cell exhaustion mediators [46]. Interestingly, correlations between tumor size and TIL density were not only non-significant but also negative in the combined durvalumab and oleclumab treatment group. Adenosine is a key inducer of vasodilation in hypoxic environments, such as within a solid tumor, and is critical in conditions like myocardial ischemia [47]. In the present study, oleclumab monotherapy resulted in the lowest median intratumoral density of total human leukocytes, implying that oleclumab-mediated inhibition of adenosine generation may have impaired leukocyte extravasation into tumor tissue via induction of intra- and peritumoral vasoconstriction. This would restrict the extent of anti-tumor immune response in the durvalumab-oleclumab treatment group to the leukocytes already present in the TME. Durvalumab’s inability to overcome Treg-mediated immunosuppression could have consequently led to the deceleration of mutually regulated CD8^+^ and FoxP3^+^ TIL proliferation, allowing tumor growth to surpass TIL expansion, resulting in diminished TIL density.

Analyses of the abundances and distribution of T-cell subsets in the peripheral blood of the experimental mice revealed accumulation of central memory CD8^+^ T cells, alongside a notable absence of effector memory and effector CD8^+^ T cells across all treatment groups. We observed a similar association between durvalumab-oleclumab treatment failure and a lack of circulating effector CD8^+^ T cells in our previous study profiling peripheral blood of NSGO-OV-UMB1/ENGOT-OV30 patients. While the majority of non-responding patients in the study exhibited a peripheral blood CD8^+^ T-cell compartment composition reflective of weak effector potential, the only responder in the clinical trial displayed a strikingly high abundance of peripheral blood effector CD8^+^ T cells, several-fold greater than in non-responders [14]. The observations of both studies suggest that the T cells in the mice, while capable of recognizing and infiltrating HGSOC, were unable to achieve full activation or maintain their functional state. Furthermore, Tregs are able to prevent effector T-cell activation and regeneration while maintaining central memory T-cell abundance [48]. Our observation of a consistently low CD8^+^ T-cell/Treg ratio among treatment groups (Fig. 3C) is in agreement with Geels et al.’s study on PD-1/PD-L1 interaction inhibition [46] and strongly implicates Tregs as a primary driver of immunosuppression in our model.

While the consistency in the development of an immune-excluded immunophenotype and comprehensive immunotherapy resistance underscore the stability of our model, as well as its faithful portrayal of the adaptive immunosuppression within the HGSOC TME, it also accentuates a limitation of our study regarding the lack of diversity within the study cohort. The use of a single cord blood donor and a single tumor material donor effectively restricts this study to a preclinical trial on a single patient, limiting our model’s generalizability to the broader HGSOC patient population. Moreover, had this approach been incorporated into the decision-making process for establishing a clinical trial, the responsiveness of the sole engrafted tumor material to immunotherapy would have been the exclusive determinant of its approval. Future research will therefore focus on developing a diverse preclinical model portfolio incorporating HSCs and HGSOC tumors from various donors, capturing different patient characteristics, disease stages, and immunophenotypes. Our model’s value could be further enhanced by using tumor material and bone marrow HSCs from the same patient to construct more personalized “avatar” models.

## 5 Conclusion

This proof-of-concept study introduces a humanized orthotopic PDX mouse model of HGSOC that effectively replicates the morphology, immune contexture, and immunotherapy resistance of the immunosuppressive HGSOC TME. We demonstrate the model’s translational potential and feasibility in analyses of immune suppression mechanisms and tumor-immune interactions within the TME of HGSOC. Our model offers a platform for preclinical evaluation of combination immunotherapies in HGSOC, enhancing the potential for successful translation of promising preclinical results to clinical trials and improvements in patient care.

## Abbreviations

BLI: bioluminescence imaging
FBS: fetal bovine serum
FIGO: International Federation of Gynecology and Obstetrics
GFP: green fluorescent protein
HGSOC: high-grade serous ovarian cancer
HSC: hematopoietic stem cell
ICI: immune checkpoint inhibitor
ICOS: inducible costimulator
IHC: immunohistochemistry
IM: invasive margin
MACS: magnetic activated cell sorting
MNC: mononuclear cell
NSG: *NOD.Cg-Prkdcscid Il2rgtm1Wj/SzJ*
NSGS: *NOD.Cg-Prkdcscid Il2rgtm1Wjl Tg (CMV-IL3, CSF2, KITLG) 1Eav/MloySzJ*
PBS: phosphate-buffered saline
PDX: patient-derived xenograft
RBC: red blood cell
RT: room temperature
TAM: tumor-associated macrophage
TIL: tumor-infiltrating leukocyte
TME: tumor microenvironment
Treg: regulatory T cell

## Resource Availability

### Lead Contact

Requests for further information and resources should be directed to and will be fulfilled by the lead contact, Dr. Katrin Kleinmanns, PhD.

### Materials Availability

This study did not generate new unique reagents.

### Data and Code Availability

All data reported in this paper will be shared by the lead contact upon request. This paper does not report original code.

Any additional information required to reanalyze the data reported in this paper is available from the lead contact upon request.

## Acknowledgements

We thank the patients for their consent and participation in the study. We thank Brith Bergum and Jørn Skavland at the Flow Cytometry Core Facility of the University of Bergen for providing support for our flow cytometry work, as well as Hege Avsnes Dale and Endy Spriet at the Molecular Imaging Center of the University of Bergen for their assistance with tissue imaging. We also declare the use of the BioRender.com platform for figure creation. This research was funded by the Western Norway Regional Health Authority (project number 28543) and the Research Council of Norway through its Centers of Excellence funding scheme (project number 223250).

## CRediT Authorship Contribution Statement

**Luka Tandaric:** Conceptualization, Methodology, Formal Analysis, Investigation, Resources, Data Curation, Writing - Original Draft, Writing - Review & Editing, Visualization; **Line Bjørge:** Conceptualization, Methodology, Investigation, Resources, Writing - Original Draft, Writing - Review & Editing, Supervision, Project Administration, Funding Acquisition; **Martine Rott Lode:** Formal Analysis, Investigation, Data Curation, Writing - Review & Editing; **Cecilie Fredvik Torkildsen:** Methodology, Investigation, Resources, Writing - Review & Editing; **Pia Aehnlich:** Methodology, Formal Analysis, Investigation, Resources, Data Curation, Writing - Review & Editing, Visualization; **Rammah Elnour:** Methodology, Resources, Data Curation, Writing - Review & Editing; **Daniela Elena Costea:** Methodology, Formal Analysis, Investigation, Resources, Writing - Review & Editing; **Lars Andreas Akslen:** Resources, Writing - Review & Editing, Supervision, Funding Acquisition; **Liv Cecilie Vestrheim Thomsen:** Writing - Original Draft, Writing - Review & Editing, Supervision; **Emmet Mc Cormack:** Resources, Writing - Review & Editing, Supervision, Funding Acquisition; **Katrin Kleinmanns:** Conceptualization, Methodology, Formal Analysis, Investigation, Resources, Data Curation, Writing - Original Draft, Writing - Review & Editing, Visualization, Supervision, Project Administration

---

**Fig. S1.**
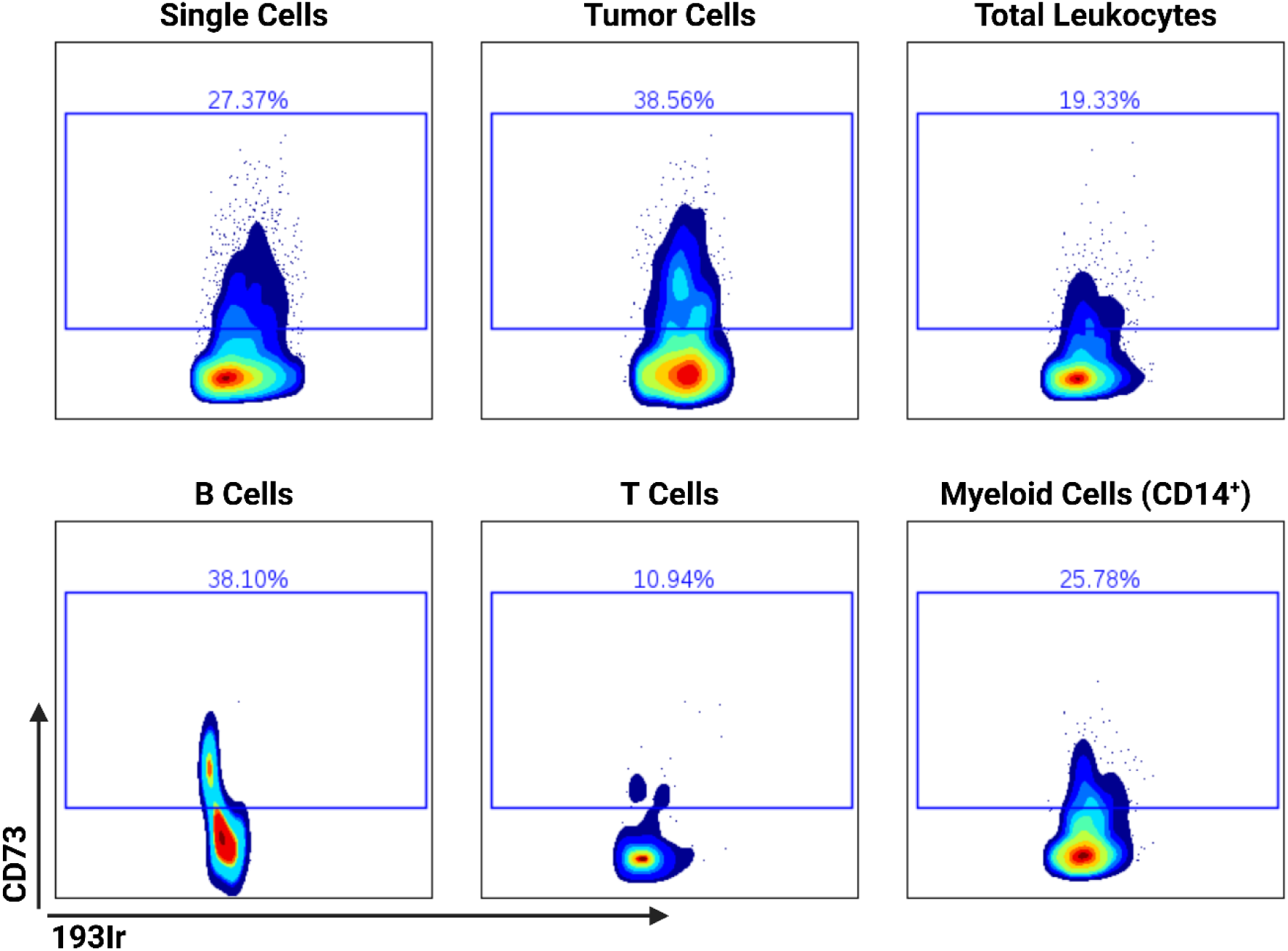
CD73 expression profiles of the constituents of the dissociated PDX material used in this study. Gates encompass cells above the CD73 positivity threshold, with the relative abundance of CD73-positive cells within the specified cell population displayed on top of each gate. The single-cell data used for this analysis was previously acquired using suspension mass cytometry.

**Fig. S2.**
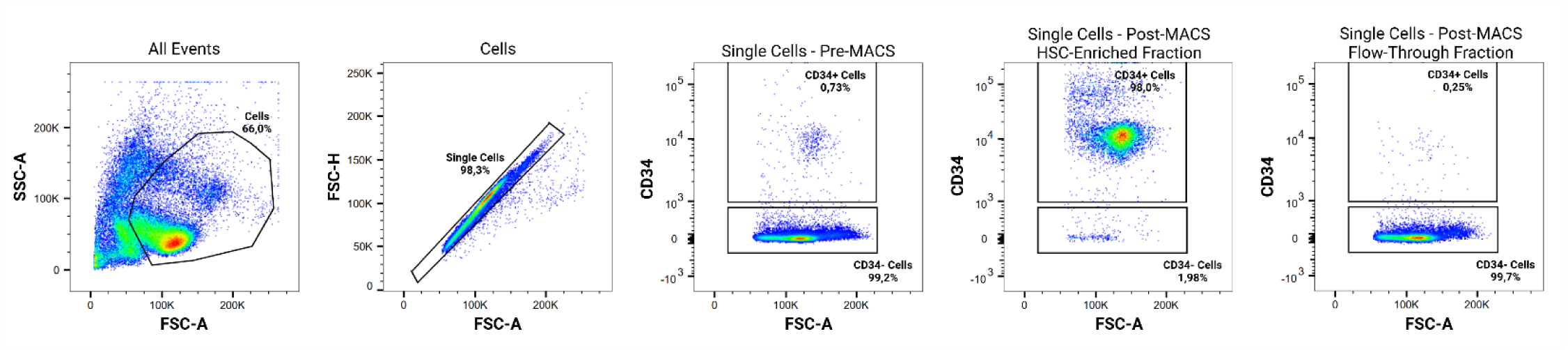
Gating strategy used for assessing the purity of samples enriched with human CD34^+^ hematopoietic stem cells from umbilical cord blood prior to their intravenous injection into NSGS mice. Enrichment was performed by magnetic activated cell sorting. Samples were analyzed using conventional flow cytometry. SSC - side scatter; FSC - forward scatter; A - area; H - height; HSC - hematopoietic stem cell; MACS - magnetic activated cell sorting

**Fig. S3.**
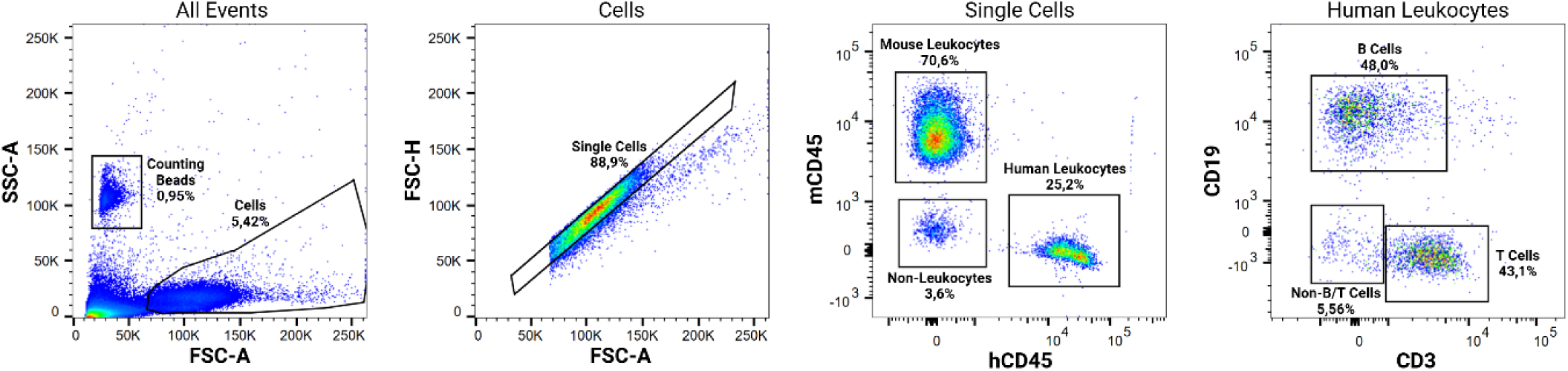
Representative gating strategy for the assessment of blood chimerism in mice injected with human hematopoietic stem cells. Leukocyte phenotyping was performed using fluorescence flow cytometry. Complete blood chimerism data is available in Table S4. SSC - side scatter; FSC - forward scatter; A - area; H - height; mCD45 - murine CD45; hCD45 - human CD45

**Fig. S4.**
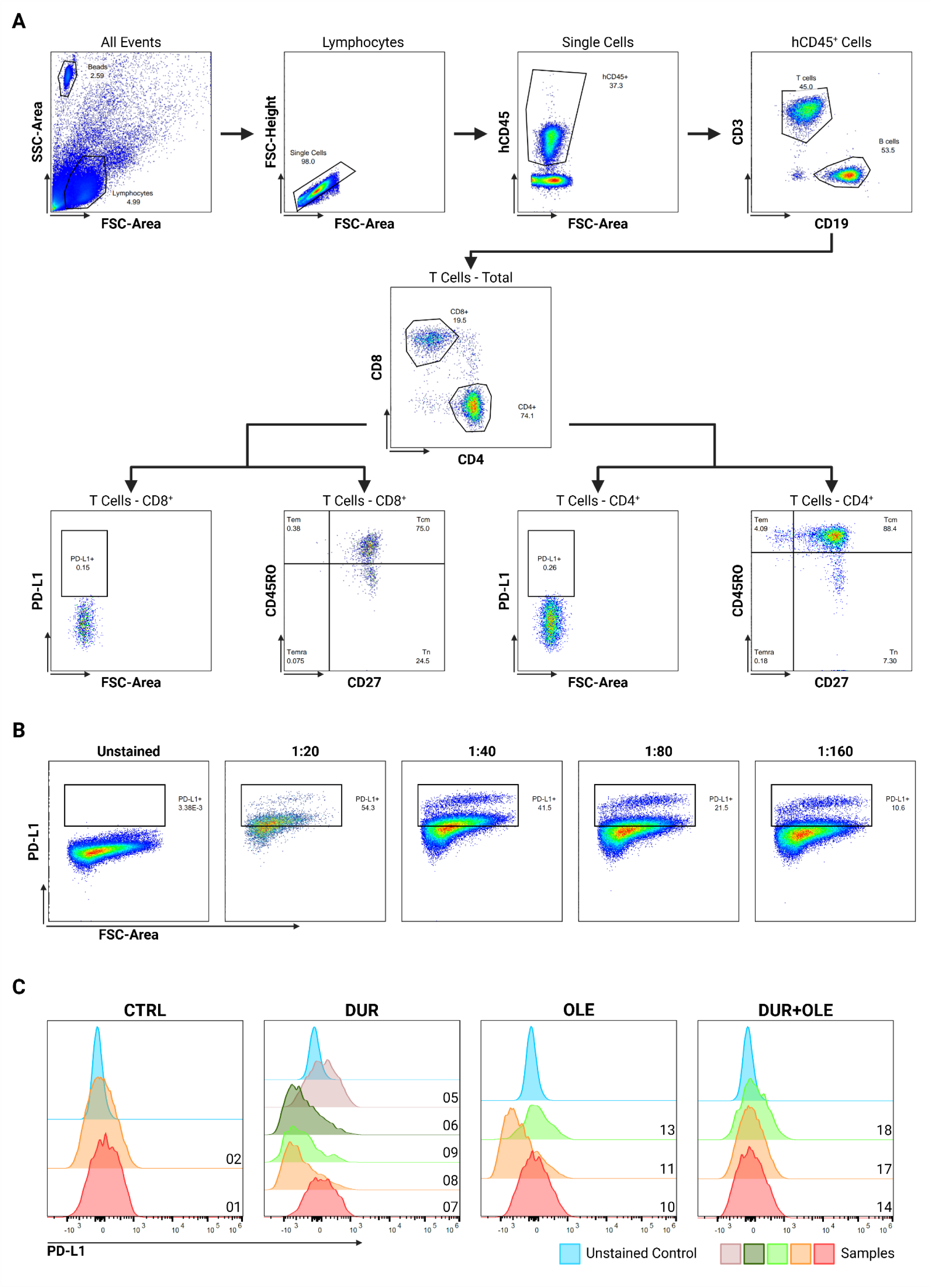
Key elements of the workflow used for the analysis of endpoint blood samples by spectral flow cytometry. (A) Titration of the anti-PD-L1 antibody. The sample used for titration consisted of peripheral blood mononuclear cells stimulated with phytohemagglutinin. The remainder of the spectral flow cytometry panel was titrated prior to this study (unpublished data). (B) Representative gating strategy used for the characterization of blood sample composition. (C) Histograms representing PD-L1 expression profiles of the total T-cell populations relative to the unstained control. Mouse IDs are displayed to the right of each histogram. Certain samples were excluded from this analysis due to low T-cell counts. Full PD-L1 expression data is available in Table S13. SSC - side scatter; FSC - forward scatter; Tn - naïve T cells; Tcm - central memory T cells; Tem - effector memory T cells; Temra - effector T cells; CTRL - control group; DUR - durvalumab-only group; OLE - oleclumab-only group; DUR+OLE - combination treatment group

**Fig. S5.**
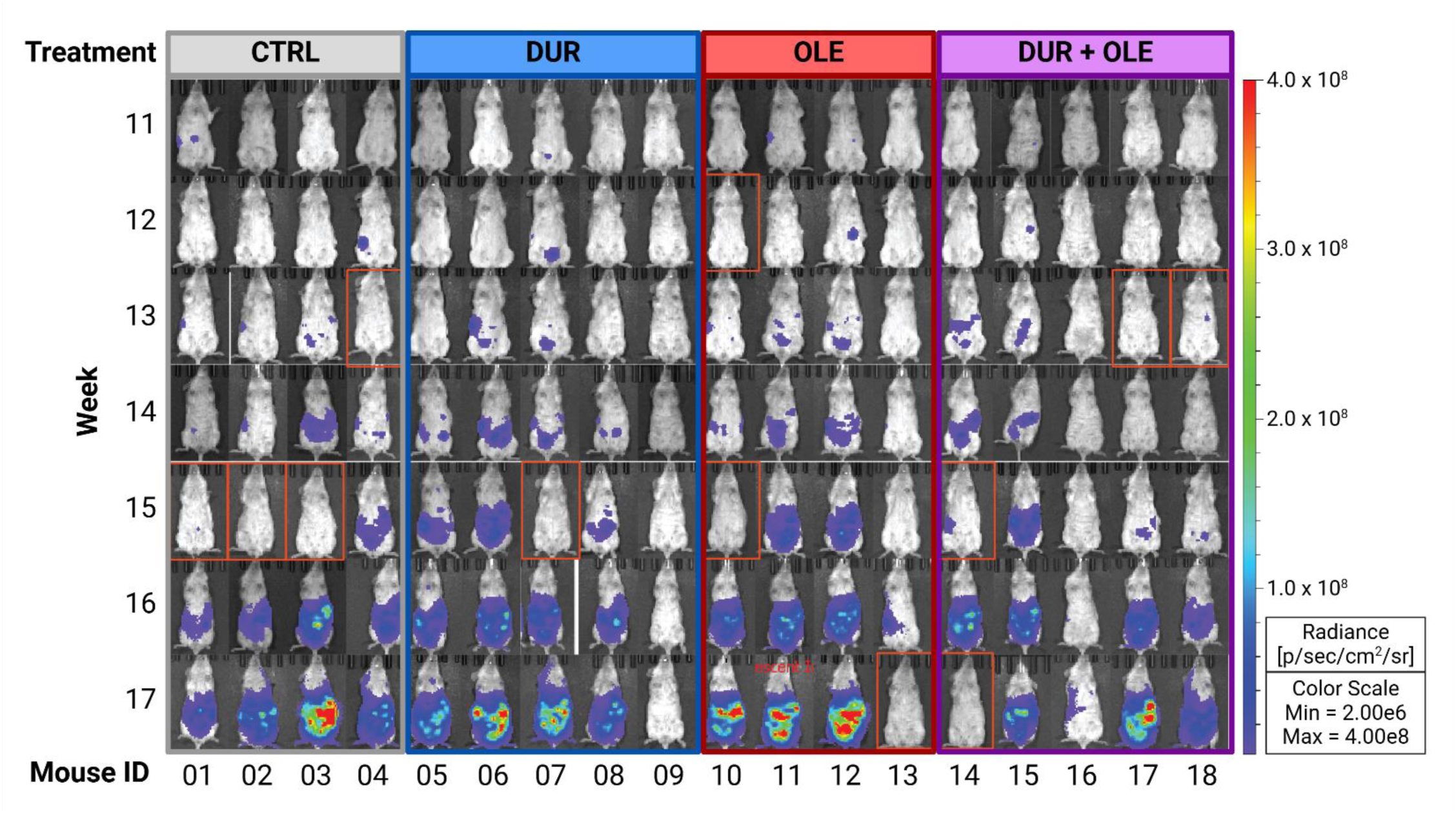
Longitudinal overview of weekly bioluminescence imaging results for PDX-implanted experimental mice in the ventral position. Each column represents an individual mouse. Only images from baseline (week 11) through endpoint (week 17) are shown. Mice outlined with an orange border displayed inexplicably low bioluminescence at the specified timepoint, even after luciferin re-injection. These data were excluded from further analyses. Full data on the total ventral flux are available in Table S5. CTRL - control group; DUR - durvalumab-only group; OLE - oleclumab-only group; DUR+OLE - combination treatment group

**Fig. S6.**
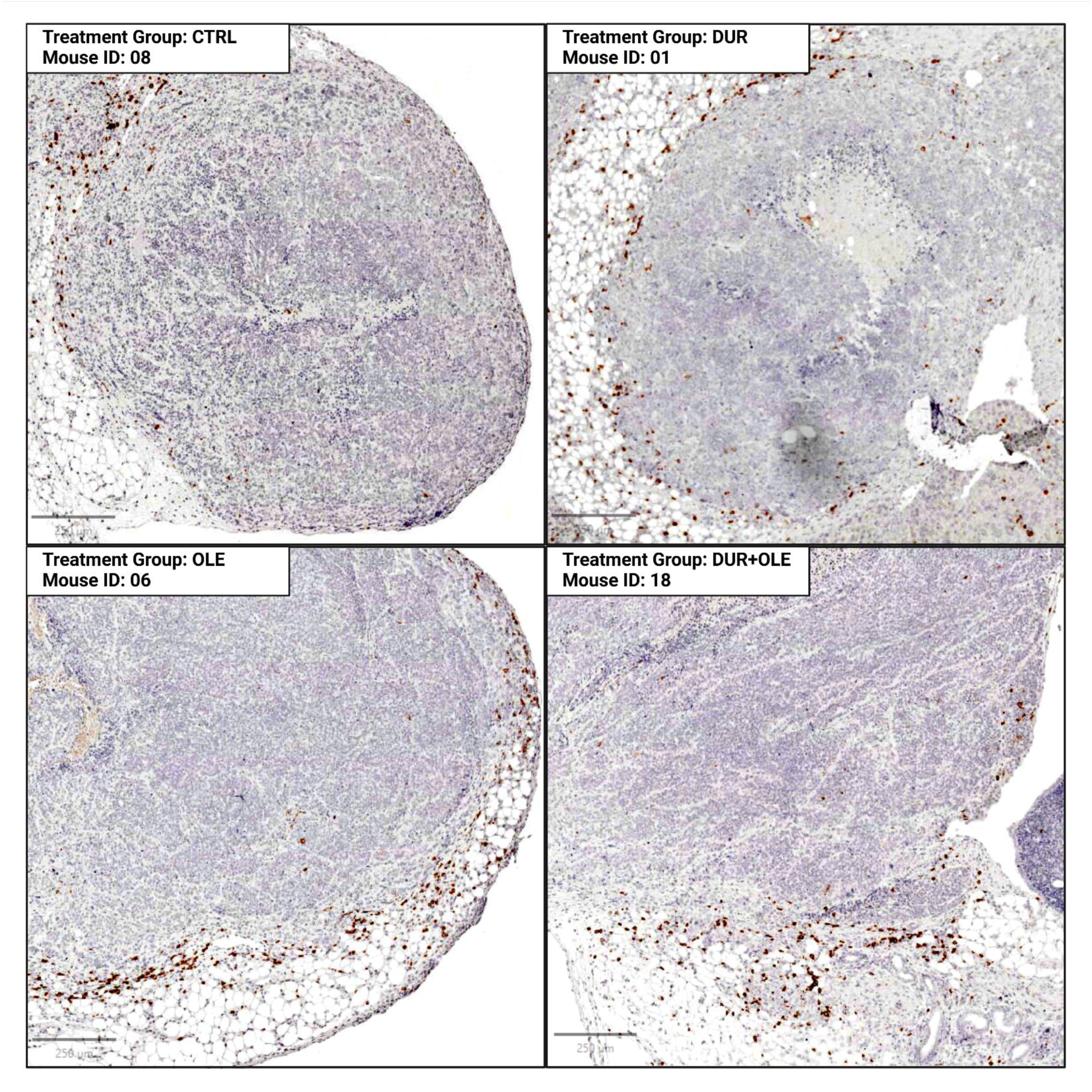
Light microscopy images (400x) of representative areas of the primary PDX tumor displaying prominent accumulation of human leukocytes in the invasive margin. Tumor sections were stained for human CD45. Each image encompasses a 2.5 mm^2^ area of a representative PDX tumor section from each treatment group (specified in the top left corner of each image). CTRL - control group; DUR - durvalumab-only group; OLE - oleclumab-only group; DUR+OLE - combination treatment group; PDX - patient-derived xenograft

**Fig. S7.**
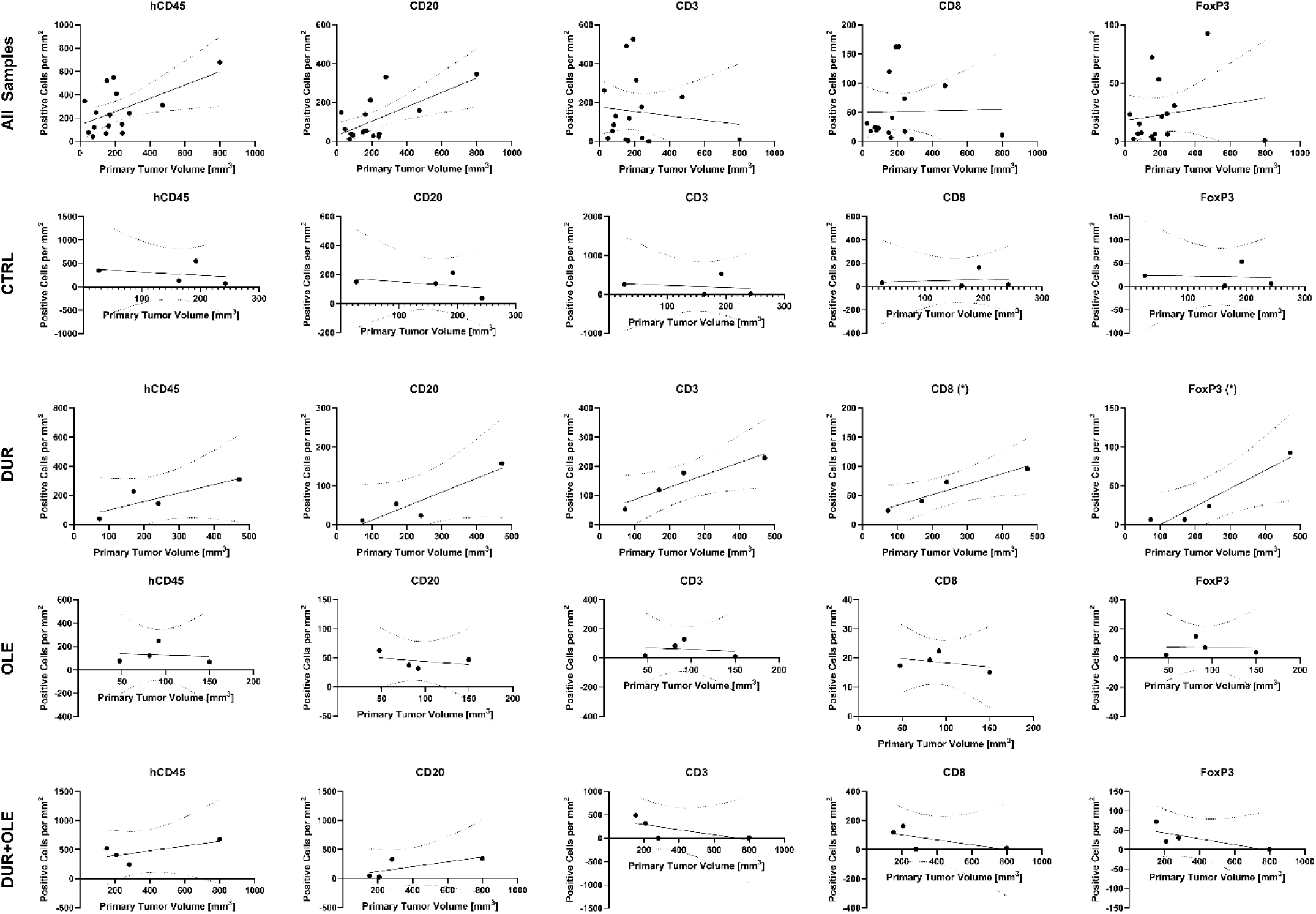
Correlation plots showing associations between tumor burden at the end of the study and densities of intratumoral marker-positive leukocytes. Top row: all samples combined. Bottom four rows: samples from individual treatment groups. Significant correlations (p<0.05) are marked with an asterisk (*) in the graph title. Full correlation data is available in Table S9. CTRL - control group; DUR - durvalumab-only group; OLE - oleclumab-only group; DUR+OLE - combination treatment group

**Table S1.** Antibody panel used for the characterization of leukocytes in the blood samples from the experimental mice using spectral flow cytometry.

| Fluorophore | Target | Clone | Dilution | Vendor | Cat.No. |
| --- | --- | --- | --- | --- | --- |
| BUV615 | CD3 | HIT3a | 1:160 | BD | 751157 |
| PerCP/Fire 806 | CD4 | SK3 | 1:160 | BioLegend | 344694 |
| Spark Blue 550 | CD8 | SK1 | 1:160 | BioLegend | 344760 |
| PE/Fire 640 | CD19 | HIB19 | 1:160 | BioLegend | 302274 |
| Spark Red 718 | CD27 | QA17A18 | 1:80 | BioLegend | 393218 |
| BV480 | CD45RO | UCHL1 | 1:160 | BD | 566143 |
| BUV 805 | CD45 | HI30 | 1:160 | ThermoFisher | 368-0459-42 |
| BV711 | PD-L1 | 29E.2A3 | 1:80 | BioLegend | 329722 |

BD - Becton, Dickinson and Company; BUV - Brilliant Ultra Violet; BV - Brilliant Violet

**Table S2.** List of antibodies used for the immunohistochemical staining of primary patient-derived xenograft tumor sections.

| Target | Clone | Host Species | Dilution | Vendor | Cat.No. |
| --- | --- | --- | --- | --- | --- |
| hCD45 | 2B11 | Mouse | 1:100 | ThermoFisher | 14-9457-82 |
| CD20 | H1 | Mouse | 1:200 | BD | 555677 |
| CD3 | F7.2.38 | Mouse | 1:200 | Abcam | Ab17143 |
| CD8 | C8/144B | Mouse | 1:400 | BioLegend | 372902 |
| FoxP3 | 236A/E7 | Mouse | 1:50 | ThermoFisher | 14-4777-82 |

BD - Becton, Dickinson and Company; hCD45 - Human CD45

**Table S3.** Overview of the positive cell detection parameters used for the enumeration of leukocytes in primary PDX tumor sections.

| Parameter | hCD45, CD3, CD8 | CD20 | FoxP3 |
| --- | --- | --- | --- |
| Detection image | Optical density sum |  |  |
| Requested pixel size | 0.17 $\mu\text{m}$ | | |
| Background Radius | 8 $\mu\text{m}$ | | |
| Use opening by reconstruction | Yes |  |  |
| Median filter radius | 0 |  |  |
| Sigma | 1.5 |  |  |
| Minimum area | 20 |  |  |
| Maximum area | 201 |  |  |
| Threshold | 0.1 |  |  |
| Max background intensity | 2 |  |  |
| Split by shape | Yes |  |  |
| Exclude DAB (membrane staining) | No |  |  |
| Cell expansion | 3 $\mu\text{m}$ | | |
| Include cell nucleus | Yes |  |  |
| Smooth boundaries | No |  |  |
| Make measurements | Yes |  |  |
| Score compartment | Cell: DAB OD Mean | Nucleus: DAB OD Mean | Nucleus: DAB OD Mean |
| Threshold 1+ | 0.04 | 0.2 | 0.3 |
| Threshold 2+ | 0.4 |  |  |
| Threshold 3+ | 0.6 |  |  |
| Single threshold | Yes | Yes | Yes |

**Table S4.** Results of the chimerism assessment of mouse blood during model establishment and at the end of the study.

| Mouse ID | Week 8<br>(PDX Implantation) |  |  | Week 11<br>(Group Assignment) |  |  | Week 17<br>(End of Study) |  |  |
| --- | --- | --- | --- | --- | --- | --- | --- | --- | --- |
|  | Human<sup>a</sup> | B Cells<sup>b</sup> | T Cells<sup>b</sup> | Human<sup>a</sup> | B Cells<sup>b</sup> | T Cells<sup>b</sup> | Human<sup>a</sup> | B Cells<sup>b</sup> | T Cells<sup>b</sup> |
| 01 | 18.4 % | 90.7 % | N/A | 19.0 % | 88.8 % | 1.43 % | 21.9 % | 64.3 % | 28.8 % |
| 02 | 18.4 % | 88.5 % | N/A | 22.6 % | 85.3 % | 1.96 % | 51.6 % | 39.7 % | 51.3 % |
| 03 | 19.4 % | 89.1 % | N/A | 19.3 % | 88.4 % | 1.11 % | 24.0 % | 91.0 % | 0.75 % |
| 04 | 17.5 % | 86.6 % | N/A | 21.1 % | 86.4 % | 1.21 % | 28.2 % | 90.5 % | 0.96 % |
| 05 | 30.9 % | 87.4 % | N/A | 37.0 % | 88.8 % | 0.75 % | 22.6 % | 47.9 % | 42.7 % |
| 06 | 18.4 % | 86.6 % | N/A | 19.2 % | 82.4 % | 2.35 % | 29.1 % | 56.0 % | 31.7 % |
| 07 | 16.9 % | 86.5 % | N/A | 24.3 % | 86.9 % | 1.66 % | 20.0 % | 41.8 % | 48.1 % |
| 08 | 17.7 % | 87.0 % | N/A | 18.6 % | 56.2 % | 30.7 % | 22.2 % | 10.7 % | 77.3 % |
| 09 | 14.2 % | 87.1 % | N/A | 13.3 % | 86.8 % | 1.61 % | 11.9 % | 78.1 % | 14.9 % |
| 10 | 13.4 % | 85.6 % | N/A | 26.3 % | 88.3 % | 1.17 % | 28.7 % | 44.5 % | 41.6 % |
| 11 | 14.1 % | 90.8 % | N/A | 23.0 % | 88.8 % | 1.10 % | 29.4 % | 32.2 % | 57.4 % |
| 12 | 14.0 % | 88.2 % | N/A | 17.9 % | 87.0 % | 1.52 % | 25.8 % | 88.7 % | 0.68 % |
| 13 | 27.6 % | 88.2 % | N/A | 28.2 % | 87.0 % | 0.91 % | 29.8 % | 87.9 % | 1.52 % |
| 14 | 12.1 % | 75.0 % | N/A | 28.8 % | 89.6 % | 1.30 % | 39.4 % | 12.2 % | 79.6 % |
| 15 | 26.6 % | 88.9 % | N/A | 31.8 % | 87.5 % | 1.29 % | 20.0 % | 75.3 % | 2.12 % |
| 16 | 13.1 % | 92.6 % | N/A | 10.6 % | 90.9 % | 0.90 % | 8.14 % | 87.6 % | 1.42 % |
| 17 | 20.9 % | 90.4 % | N/A | 19.8 % | 80.3 % | 6.66 % | 30.4 % | 22.9 % | 70.6 % |
| 18 | 13.3 % | 88.6 % | N/A | 18.2 % | 86.8 % | 1.16 % | 14.2 % | 82.6 % | 9.52 % |

<sup>a</sup> Percentage of single cells that are positive for human CD45.
<sup>b</sup> Relative to total cells positive for human CD45.

**Table S5.** Total lateral and ventral photon flux measured weekly during weeks 9 through 17 after injection of hematopoietic cells into the experimental mice.

| Mouse ID | W09 | W10 | W11 | W12 | W13 | W14 | W15 | W16 | W17 |
| --- | --- | --- | --- | --- | --- | --- | --- | --- | --- |
| **Total Flux - Lateral** |  |  |  |  |  |  |  |  |  |
| 01 | 1.49E+07 | 6.66E+07 | 2.67E+08 | 1.74E+09 | 2.43E+09 | 3.25E+09 | 5.00E+07 | 7.69E+09 | 5.98E+09 |
| 02 | 1.03E+07 | 2.74E+07 | 3.86E+08 | 7.00E+08 | 1.42E+09 | 3.38E+09 | 1.90E+06 | 3.49E+09 | 1.11E+10 |
| 03 | 3.93E+05 | 7.23E+07 | 2.67E+07 | 1.22E+09 | 2.87E+09 | 4.27E+09 | 3.76E+06 | 1.00E+10 | 2.44E+10 |
| 04 | 4.40E+05 | 4.84E+07 | 4.25E+08 | 1.32E+09 | 1.72E+06 | 5.85E+09 | 6.78E+09 | 1.11E+10 | 1.24E+10 |
| 05 | 4.39E+06 | 1.52E+07 | 8.25E+07 | 2.02E+08 | 1.40E+09 | 2.19E+09 | 3.30E+09 | 5.18E+09 | 6.60E+09 |
| 06 | 2.43E+07 | 9.86E+07 | 7.16E+08 | 2.92E+06 | 8.37E+09 | 6.81E+09 | 6.05E+09 | 1.29E+10 | 2.26E+10 |
| 07 | 4.43E+05 | 3.69E+07 | 2.36E+08 | 5.47E+08 | 1.48E+09 | 2.73E+09 | 4.21E+07 | 8.16E+09 | 1.45E+10 |
| 08 | 2.32E+06 | 8.93E+06 | 1.00E+08 | 8.16E+07 | 2.19E+08 | 1.53E+09 | 4.48E+08 | 3.84E+09 | 4.88E+09 |
| 09 | 7.80E+05 | 8.69E+05 | 2.30E+06 | 1.72E+06 | 3.82E+05 | 2.87E+07 | 5.17E+07 | 1.59E+06 | 2.95E+06 |
| 10 | 1.37E+06 | 6.96E+07 | 2.98E+08 | 2.93E+06 | 5.05E+09 | 5.81E+09 | 1.11E+08 | 1.10E+10 | 1.93E+10 |
| 11 | 1.36E+07 | 7.39E+07 | 2.35E+08 | 1.13E+09 | 2.65E+09 | 2.37E+09 | 4.00E+09 | 1.25E+10 | 2.44E+10 |
| 12 | 6.18E+05 | 1.63E+08 | 6.25E+08 | 2.42E+09 | 6.67E+09 | 1.36E+10 | 1.17E+10 | 1.56E+10 | 4.46E+10 |
| 13 | 4.43E+06 | 1.63E+07 | 4.49E+07 | 2.29E+08 | 5.03E+08 | 1.71E+09 | 2.92E+09 | 5.52E+09 | 1.26E+07 |
| 14 | 2.38E+07 | 6.75E+07 | 4.28E+08 | 2.03E+09 | 4.49E+09 | 5.52E+09 | 2.80E+07 | 1.43E+10 | 1.53E+07 |
| 15 | 7.69E+06 | 5.18E+07 | 3.39E+08 | 1.01E+09 | 2.69E+09 | 4.01E+09 | 5.42E+09 | 7.79E+09 | 7.86E+09 |
| 16 | 1.23E+06 | 6.33E+06 | 4.67E+07 | 7.20E+07 | 7.39E+08 | 7.30E+08 | 1.33E+09 | 5.80E+08 | 1.70E+09 |
| 17 | 5.89E+05 | 1.87E+07 | 2.22E+08 | 3.53E+08 | 6.41E+05 | 2.26E+07 | 5.51E+08 | 1.08E+10 | 2.11E+10 |
| 18 | 1.00E+06 | 1.68E+06 | 1.43E+07 | 4.19E+07 | 9.87E+05 | 2.22E+08 | 1.27E+09 | 1.90E+09 | 5.15E+09 |
| **Total Flux - Ventral** |  |  |  |  |  |  |  |  |  |
| 01 | 2.01E+09 | 2.30E+09 | 1.13E+11 | 4.29E+10 | 8.02E+10 | 1.07E+11 | 5.55E+10 | 8.12E+11 | 2.69E+12 |
| 02 | 9.34E+08 | 7.25E+08 | 1.86E+09 | 1.70E+10 | 6.68E+10 | 1.39E+11 | 1.07E+09 | 1.79E+12 | 1.02E+13 |
| 03 | 4.78E+08 | 1.13E+09 | 1.31E+09 | 2.58E+10 | 2.13E+11 | 8.28E+11 | 2.02E+09 | 1.01E+13 | 6.11E+13 |
| 04 | 4.71E+08 | 2.74E+09 | 2.96E+09 | 3.30E+11 | 6.16E+09 | 2.27E+11 | 8.80E+11 | 2.97E+12 | 6.97E+12 |
| 05 | 8.04E+08 | 1.23E+09 | 3.48E+09 | 4.16E+09 | 6.82E+10 | 4.03E+11 | 1.49E+12 | 3.75E+12 | 1.26E+13 |
| 06 | 1.14E+09 | 1.41E+09 | 1.07E+10 | 8.71E+08 | 5.27E+11 | 1.42E+12 | 2.68E+12 | 6.89E+12 | 3.04E+13 |
| 07 | 5.22E+08 | 1.03E+10 | 4.50E+10 | 3.17E+11 | 2.26E+11 | 6.15E+11 | 1.91E+10 | 4.84E+12 | 2.43E+13 |
| 08 | 9.86E+08 | 1.28E+09 | 2.32E+09 | 9.14E+06 | 1.05E+10 | 2.06E+11 | 5.00E+11 | 3.31E+12 | 7.14E+12 |
| 09 | 1.08E+09 | 1.07E+09 | 1.15E+09 | 6.89E+08 | 1.00E+11 | 1.80E+09 | 2.96E+09 | 5.99E+08 | 6.62E+08 |
| 10 | 6.93E+08 | 1.65E+09 | 3.16E+09 | 5.98E+08 | 1.67E+11 | 2.41E+11 | 3.56E+10 | 4.99E+12 | 2.67E+13 |
| 11 | 1.34E+09 | 2.28E+09 | 8.80E+10 | 4.50E+10 | 2.44E+11 | 5.81E+11 | 2.63E+12 | 7.91E+12 | 4.05E+13 |
| 12 | 5.85E+08 | 6.96E+09 | 2.80E+10 | 2.35E+11 | 2.83E+11 | 6.34E+11 | 2.73E+12 | 7.68E+12 | 5.69E+13 |
| 13 | 1.49E+09 | 1.59E+09 | 2.02E+09 | 6.24E+09 | 3.38E+09 | 4.48E+10 | 2.60E+10 | 6.71E+11 | 1.27E+09 |
| 14 | 9.20E+08 | 2.17E+09 | 1.31E+10 | 5.20E+10 | 3.13E+11 | 7.87E+11 | 2.04E+11 | 9.18E+12 | 6.15E+09 |
| 15 | 1.54E+09 | 5.09E+09 | 1.90E+10 | 8.57E+10 | 2.88E+11 | 4.92E+11 | 2.63E+12 | 6.89E+12 | 8.13E+12 |
| 16 | 1.07E+09 | 8.63E+08 | 1.86E+09 | 6.00E+09 | 2.66E+10 | 1.57E+10 | 7.75E+10 | 6.85E+10 | 3.30E+11 |
| 17 | 6.70E+08 | 2.98E+09 | 1.81E+09 | 9.92E+09 | 3.73E+08 | 5.62E+09 | 1.94E+11 | 4.68E+12 | 3.19E+13 |
| 18 | 1.04E+09 | 9.25E+08 | 1.65E+09 | 1.44E+09 | 5.75E+08 | 1.87E+10 | 1.57E+11 | 1.15E+12 | 4.52E+12 |

W - week

**Table S6.** Dimensions of primary patient-derived xenograft tumors measured at the end of the study. Tumor volumes were calculated using the formula: (height x width x length x π) / 6.

| Mouse ID | Tumor Dimensions |  |  |  |
| --- | --- | --- | --- | --- |
|  | Height [mm] | Width [mm] | Length [mm] | Volume [mm<sup>3</sup>] |
| 01 | 2.8 | 3.4 | 5.32 | 26.52 |
| 02 | 8.35 | 3.32 | 13.24 | 192.18 |
| 03 | 5.73 | 8.38 | 9.65 | 242.62 |
| 04 | 6.47 | 6.1 | 7.88 | 162.84 |
| 05 | 11.27 | 7.45 | 10.73 | 471.71 |
| 06 | 4.99 | 6.14 | 10.61 | 170.21 |
| 07 | 4.34 | 3.26 | 6.72 | 49.78 |
| 08 | 5.8 | 9.39 | 8.43 | 240.39 |
| 09 | 4.2 | 5.04 | 6.57 | 72.82 |
| 10 | 6.92 | 5.51 | 4.07 | 81.26 |
| 11 | 5.03 | 6.54 | 5.33 | 91.81 |
| 12 | 5.43 | 7.66 | 6.88 | 149.84 |
| 13 | 5.17 | 4.86 | 3.59 | 47.23 |
| 14 | 7.4 | 4.91 | 8.04 | 152.96 |
| 15 | 10.09 | 10.72 | 14.11 | 799.12 |
| 16 | 10.75 | 7.02 | 7.13 | 281.73 |
| 17 | 6.2 | 10.8 | 5.95 | 208.61 |
| 18<sup>a</sup> | - | - | - | - |

<sup>a</sup> Primary tumor too small to measure.

**Table S7.** The extent of visible metastatic dissemination at the end of the study.

| Mouse ID | Anatomic Localization of Visible Metastatic Lesion(s) |  |  |  |  |  |  |  |  |
| --- | --- | --- | --- | --- | --- | --- | --- | --- | --- |
|  | DIA | DUO | GAS | HEP | MES | OME | PER | REN | SPL |
| 01 |  |  |  | X |  |  | X |  | X |
| 02 |  |  |  | X |  |  | X |  |  |
| 03 | X |  | X |  |  |  |  |  |  |
| 04 |  |  |  |  |  | X | X |  |  |
| 05 |  |  |  |  | X | X |  | X |  |
| 06 | X |  |  |  |  |  | X |  |  |
| 07 | X |  |  |  |  |  | X | X | X |
| 08 | X | X |  |  |  | X | X |  |  |
| 09 |  |  |  |  |  |  |  |  |  |
| 10 |  |  |  |  |  | X | X |  |  |
| 11 |  |  |  |  |  |  | X |  |  |
| 12 | X |  |  | X |  | X | X |  |  |
| 13 |  |  |  |  |  |  | X |  |  |
| 14 | X |  |  |  |  | X | X |  |  |
| 15 | X |  | X |  |  | X | X | X | X |
| 16 | X |  |  |  |  |  |  |  |  |
| 17 |  |  |  |  |  |  |  |  |  |
| 18 | X |  |  |  |  |  |  |  |  |

DIA - diaphragmatic; DUO - duodenal; GAS - gastric; HEP - hepatic; MES - mesenteric; OME - omental; PER - peritoneal wall; REN - renal; SPL - splenic

**Table S8.** Results of the digital analysis of primary patient-derived xenograft tumor sections. The total size of the annotated area is expressed as mm^2^, and the positive cell density is expressed as the total number of positive cells per mm^2^ of the total annotated area.

| Mouse ID | Parameter | Marker |  |  |  |  |
| --- | --- | --- | --- | --- | --- | --- |
|  |  | hCD45 | CD20 | CD3 | CD8 | FoxP3 |
| 01 | Area | 0.65 | 0.65 | 0.65 | 0.51 | 0.65 |
|  | Count | 226 | 97 | 171 | 16 | 15 |
|  | Density | 346.4 | 148.7 | 262.1 | 31.1 | 23.2 |
| 02 | Area | 1.99 | 1.99 | 1.97 | 1.99 | 1.95 |
|  | Count | 1094 | 424 | 1037 | 324 | 104 |
|  | Density | 548.5 | 212.6 | 526.1 | 162.4 | 53.3 |
| 03 | Area | 0.81 | 0.80 | 0.81 | 0.76 | 0.78 |
|  | Count | 58 | 30 | 15 | 13 | 5 |
|  | Density | 71.2 | 37.6 | 18.4 | 17.1 | 6.4 |
| 04 | Area | 1.60 | 1.60 | 1.60 | 1.50 | 1.60 |
|  | Count | 215 | 221 | 6 | 10 | 3 |
|  | Density | 134.4 | 138.2 | 3.8 | 6.7 | 1.9 |
| 05 | Area | 1.47 | 1.46 | 1.47 | 1.47 | 1.47 |
|  | Count | 459 | 230 | 338 | 141 | 136 |
|  | Density | 311.3 | 157.5 | 229.2 | 95.6 | 92.7 |
| 06 | Area | 1.38 | 1.38 | 1.37 | 1.38 | 1.37 |
|  | Count | 316 | 74 | 164 | 56 | 9 |
|  | Density | 228.8 | 53.8 | 119.6 | 40.5 | 6.5 |
| 07<sup>a</sup> | - | - | - | - | - | - |
| 08 | Area | 0.79 | 0.79 | 0.79 | 0.76 | 0.79 |
|  | Count | 116 | 19 | 141 | 56 | 19 |
|  | Density | 146.0 | 23.9 | 177.5 | 73.4 | 23.9 |
| 09 | Area | 0.90 | 0.90 | 0.80 | 0.90 | 0.90 |
|  | Count | 37 | 10 | 43 | 22 | 6 |
|  | Density | 40.9 | 11.1 | 53.4 | 24.3 | 6.7 |
| 10 | Area | 0.91 | 0.93 | 0.93 | 0.93 | 0.93 |
|  | Count | 110 | 35 | 79 | 18 | 14 |
|  | Density | 121.1 | 37.5 | 84.6 | 19.3 | 15.0 |
| 11 | Area | 1.34 | 1.29 | 1.33 | 1.34 | 1.34 |
|  | Count | 330 | 41 | 174 | 30 | 10 |
|  | Density | 247.0 | 31.8 | 130.6 | 22.5 | 7.5 |
| 12 | Area | 1.00 | 1.00 | 1.00 | 0.99 | 0.98 |
|  | Count | 68 | 47 | 9 | 15 | 4 |
|  | Density | 67.8 | 46.8 | 9.0 | 15.1 | 4.1 |
| 13 | Area | 0.86 | 0.86 | 0.85 | 0.86 | 0.86 |
|  | Count | 66 | 54 | 14 | 15 | 2 |
|  | Density | 76.7 | 62.8 | 16.5 | 17.4 | 2.3 |
| 14 | Area | 0.69 | 0.69 | 0.69 | 0.69 | 0.69 |
|  | Count | 362 | 34 | 341 | 83 | 50 |
|  | Density | 521.6 | 49.0 | 491.3 | 119.6 | 72.0 |
| 15 | Area | 1.15 | 1.15 | 0.76 | 1.06 | 1.06 |
|  | Count | 782 | 399 | 7 | 12 | 1 |
|  | Density | 679.0 | 346.5 | 9.2 | 11.3 | 0.9 |
| 16 | Area | 0.72 | 0.72 | 0.72 | 0.72 | 0.72 |
|  | Count | 174 | 238 | 1 | 3 | 22 |
|  | Density | 241.9 | 330.8 | 1.4 | 4.2 | 30.6 |
| 17 | Area | 2.53 | 2.52 | 2.53 | 2.53 | 2.43 |
|  | Count | 1034 | 69 | 795 | 411 | 51 |
|  | Density | 409.4 | 27.4 | 314.8 | 162.7 | 21.0 |
| 18 | Area | 0.86 | 0.82 | 0.86 | 0.83 | 0.86 |
|  | Count | 34 | 22 | 38 | 49 | 9 |
|  | Density | 39.4 | 26.7 | 44.0 | 59.3 | 10.4 |

<sup>a</sup> No discernable invasive tumor margin present.

**Table S9.** Parameters of the correlations between tumor burden at the end of the study and densities of intratumoral marker-positive leukocytes for all samples combined and each treatment group individually.

| Treatment Group | Parameter | Pearson's Correlation Coefficient (r) |  |  |  |  |
| --- | --- | --- | --- | --- | --- | --- |
|  |  | Goodness of Fit (R<sup>2</sup>) |  |  |  |  |
|  |  | p-value |  |  |  |  |
|  |  | hCD45 | CD20 | CD3 | CD8 | FoxP3 |
| All Samples | r<sup>a</sup> | 0.382 | 0.359 | -0.082 | -0.029 | 0.159 |
|  | R<sup>2</sup> | 0.317 | 0.433 | 0.016 | 0.001 | 0.029 |
|  | p | 0.145 | 0.173 | 0.763 | 0.917 | 0.556 |
| Control | r | -0.299 | -0.368 | -0.195 | 0.167 | -0.079 |
|  | R<sup>2</sup> | 0.089 | 0.135 | 0.038 | 0.028 | 0.006 |
|  | p | 0.701 | 0.632 | 0.805 | 0.833 | 0.921 |
| Durvalumab | r | 0.853 | 0.922 | 0.950 | 0.952 | 0.961 |
|  | R<sup>2</sup> | 0.728 | 0.850 | 0.902 | 0.907 | 0.924 |
|  | p | 0.147 | 0.078 | 0.050 | 0.048 | 0.039 |
| Oleclumab | r | -0.107 | -0.351 | -0.173 | -0.392 | -0.061 |
|  | R<sup>2</sup> | 0.012 | 0.123 | 0.030 | 0.153 | 0.004 |
|  | p | 0.893 | 0.649 | 0.828 | 0.608 | 0.939 |
| Durvalumab + Oleclumab | r | 0.659 | 0.720 | -0.680 | -0.638 | -0.756 |
|  | R<sup>2</sup> | 0.434 | 0.518 | 0.463 | 0.407 | 0.571 |
|  | p | 0.341 | 0.280 | 0.320 | 0.362 | 0.244 |

r - Pearson's correlation coefficient
R<sup>2</sup> - goodness of fit parameter of the results of simple linear regression
p - p-value
<sup>a</sup> Spearman's rank correlation used due to non-normality of dataset.

**Table S10.** Results of the spectral flow cytometry analysis of leukocytes in the blood samples from the experimental mice - human leukocyte counts.

| Mouse ID | Total Human Leukocytes | T Cells |  |  |  |  |  |  |  |  |  |  |
| --- | --- | --- | --- | --- | --- | --- | --- | --- | --- | --- | --- | --- |
|  |  | Total | CD4+ |  |  |  |  | CD8+ |  |  |  |  |
|  |  |  | Total | N | CM | EM | E | Total | N | CM | EM | E |
| 01 | 11758 | 2687 | 2281 | 115 | 1976 | 188 | 2 | 317 | 80 | 237 | 0 | 0 |
| 02 | 92523 | 47690 | 42615 | 997 | 29170 | 12211 | 237 | 4197 | 472 | 3720 | 5 | 0 |
| 03 | 20376 | 24 | 2 | 1 | 0 | 1 | 0 | 8 | 7 | 1 | 0 | 0 |
| 04 | 11458 | 18 | 0 | 0 | 0 | 0 | 0 | 1 | 1 | 0 | 0 | 0 |
| 05 | 15147 | 6823 | 5057 | 369 | 4472 | 207 | 9 | 1333 | 327 | 1000 | 5 | 1 |
| 06 | 27069 | 7650 | 4705 | 214 | 3300 | 1169 | 22 | 1583 | 154 | 1400 | 25 | 4 |
| 07 | 4735 | 2028 | 1948 | 95 | 1519 | 330 | 4 | 50 | 9 | 40 | 0 | 1 |
| 08 | 4726 | 3856 | 3344 | 30 | 2731 | 571 | 12 | 313 | 36 | 268 | 7 | 2 |
| 09 | 4616 | 345 | 257 | 67 | 121 | 65 | 4 | 48 | 27 | 21 | 0 | 0 |
| 10 | 14045 | 5025 | 4216 | 113 | 3035 | 1057 | 11 | 453 | 44 | 408 | 1 | 0 |
| 11 | 23690 | 14465 | 12695 | 252 | 10262 | 2157 | 24 | 1281 | 207 | 1069 | 3 | 2 |
| 12 | 25128 | 28 | 0 | 0 | 0 | 0 | 0 | 3 | 3 | 0 | 0 | 0 |
| 13 | 23542 | 109 | 48 | 15 | 30 | 3 | 0 | 32 | 26 | 4 | 0 | 2 |
| 14 | 16152 | 13406 | 11552 | 334 | 8703 | 2391 | 124 | 731 | 82 | 649 | 0 | 0 |
| 15 | 11661 | 28 | 14 | 4 | 10 | 0 | 0 | 6 | 5 | 1 | 0 | 0 |
| 16 | 9711 | 2 | 0 | 0 | 0 | 0 | 0 | 0 | 0 | 0 | 0 | 0 |
| 17 | 47863 | 35237 | 26539 | 1130 | 22368 | 2821 | 220 | 6140 | 297 | 5805 | 27 | 11 |
| 18 | 15658 | 1145 | 654 | 63 | 332 | 245 | 14 | 249 | 141 | 105 | 2 | 1 |

CM - Central Memory; E - Effector (Temra); EM - Effector Memory; N - Naïve

**Table S11.** Results of the spectral flow cytometry analysis of leukocytes in the blood samples from the experimental mice - human leukocyte counts per µL of blood.

| Mouse ID | Total Human Leukocytes | T Cells |  |  |  |  |  |  |  |  |  |  |
| --- | --- | --- | --- | --- | --- | --- | --- | --- | --- | --- | --- | --- |
|  |  | Total | CD4+ |  |  |  |  | CD8+ |  |  |  |  |
|  |  |  | Total | N | CM | EM | E | Total | N | CM | EM | E |
| 01 | 91.6 | 20.9 | 17.8 | 0.9 | 15.4 | 1.5 | 0.0 | 2.5 | 0.6 | 1.8 | 0.0 | 0.0 |
| 02 | 608.6 | 313.7 | 280.3 | 6.6 | 191.9 | 80.3 | 1.6 | 27.6 | 3.1 | 24.5 | 0.0 | 0.0 |
| 03 | 168.3 | 0.2 | 0.0 | 0.0 | 0.0 | 0.0 | 0.0 | 0.1 | 0.1 | 0.0 | 0.0 | 0.0 |
| 04 | 186.8 | 0.3 | 0.0 | 0.0 | 0.0 | 0.0 | 0.0 | 0.0 | 0.0 | 0.0 | 0.0 | 0.0 |
| 05 | 140.5 | 63.3 | 46.9 | 3.4 | 41.5 | 1.9 | 0.1 | 12.4 | 3.0 | 9.3 | 0.0 | 0.0 |
| 06 | 377.2 | 106.6 | 65.6 | 3.0 | 46.0 | 16.3 | 0.3 | 22.1 | 2.1 | 19.5 | 0.3 | 0.1 |
| 07 | 38.7 | 16.6 | 15.9 | 0.8 | 12.4 | 2.7 | 0.0 | 0.4 | 0.1 | 0.3 | 0.0 | 0.0 |
| 08 | 73.5 | 59.9 | 52.0 | 0.5 | 42.5 | 8.9 | 0.2 | 4.9 | 0.6 | 4.2 | 0.1 | 0.0 |
| 09 | 56.5 | 4.2 | 3.1 | 0.8 | 1.5 | 0.8 | 0.0 | 0.6 | 0.3 | 0.3 | 0.0 | 0.0 |
| 10 | 203.0 | 72.6 | 60.9 | 1.6 | 43.9 | 15.3 | 0.2 | 6.5 | 0.6 | 5.9 | 0.0 | 0.0 |
| 11 | 369.9 | 225.9 | 198.2 | 3.9 | 160.2 | 33.7 | 0.4 | 20.0 | 3.2 | 16.7 | 0.0 | 0.0 |
| 12 | 228.6 | 0.3 | 0.0 | 0.0 | 0.0 | 0.0 | 0.0 | 0.0 | 0.0 | 0.0 | 0.0 | 0.0 |
| 13 | 185.9 | 0.9 | 0.4 | 0.1 | 0.2 | 0.0 | 0.0 | 0.3 | 0.2 | 0.0 | 0.0 | 0.0 |
| 14 | 157.6 | 130.8 | 112.7 | 3.3 | 84.9 | 23.3 | 1.2 | 7.1 | 0.8 | 6.3 | 0.0 | 0.0 |
| 15 | 215.3 | 0.5 | 0.3 | 0.1 | 0.2 | 0.0 | 0.0 | 0.1 | 0.1 | 0.0 | 0.0 | 0.0 |
| 16 | 93.5 | 0.0 | 0.0 | 0.0 | 0.0 | 0.0 | 0.0 | 0.0 | 0.0 | 0.0 | 0.0 | 0.0 |
| 17 | 420.9 | 309.8 | 233.4 | 9.9 | 196.7 | 24.8 | 1.9 | 54.0 | 2.6 | 51.0 | 0.2 | 0.1 |
| 18 | 107.1 | 7.8 | 4.5 | 0.4 | 2.3 | 1.7 | 0.1 | 1.7 | 1.0 | 0.7 | 0.0 | 0.0 |

CM - Central Memory; E - Effector (Temra); EM - Effector Memory; N - Naive

**Table S12.** Results of the spectral flow cytometry analysis of leukocytes in the blood samples from the experimental mice - human leukocyte abundances relative to total human leukocytes. Values in the cells are expressed as percentages.

| Mouse ID | Total Human Leukocytes | T Cells |  |  |  |  |  |  |  |  |  |  |
| --- | --- | --- | --- | --- | --- | --- | --- | --- | --- | --- | --- | --- |
|  |  | Total | CD4+ |  |  |  |  | CD8+ |  |  |  |  |
|  |  |  | Total | N | CM | EM | E | Total | N | CM | EM | E |
| 01 | 100.0 | 22.9 | 19.4 | 1.0 | 16.8 | 1.6 | 0.0 | 2.7 | 0.7 | 2.0 | 0.0 | 0.0 |
| 02 | 100.0 | 51.5 | 46.1 | 1.1 | 31.5 | 13.2 | 0.3 | 4.5 | 0.5 | 4.0 | 0.0 | 0.0 |
| 03 | 100.0 | 0.1 | 0.0 | 0.0 | 0.0 | 0.0 | 0.0 | 0.0 | 0.0 | 0.0 | 0.0 | 0.0 |
| 04 | 100.0 | 0.2 | 0.0 | 0.0 | 0.0 | 0.0 | 0.0 | 0.0 | 0.0 | 0.0 | 0.0 | 0.0 |
| 05 | 100.0 | 45.0 | 33.4 | 2.4 | 29.5 | 1.4 | 0.1 | 8.8 | 2.2 | 6.6 | 0.0 | 0.0 |
| 06 | 100.0 | 28.3 | 17.4 | 0.8 | 12.2 | 4.3 | 0.1 | 5.8 | 0.6 | 5.2 | 0.1 | 0.0 |
| 07 | 100.0 | 42.8 | 41.1 | 2.0 | 32.1 | 7.0 | 0.1 | 1.1 | 0.2 | 0.8 | 0.0 | 0.0 |
| 08 | 100.0 | 81.6 | 70.8 | 0.6 | 57.8 | 12.1 | 0.3 | 6.6 | 0.8 | 5.7 | 0.1 | 0.0 |
| 09 | 100.0 | 7.5 | 5.6 | 1.5 | 2.6 | 1.4 | 0.1 | 1.0 | 0.6 | 0.5 | 0.0 | 0.0 |
| 10 | 100.0 | 35.8 | 30.0 | 0.8 | 21.6 | 7.5 | 0.1 | 3.2 | 0.3 | 2.9 | 0.0 | 0.0 |
| 11 | 100.0 | 61.1 | 53.6 | 1.1 | 43.3 | 9.1 | 0.1 | 5.4 | 0.9 | 4.5 | 0.0 | 0.0 |
| 12 | 100.0 | 0.1 | 0.0 | 0.0 | 0.0 | 0.0 | 0.0 | 0.0 | 0.0 | 0.0 | 0.0 | 0.0 |
| 13 | 100.0 | 0.5 | 0.2 | 0.1 | 0.1 | 0.0 | 0.0 | 0.1 | 0.1 | 0.0 | 0.0 | 0.0 |
| 14 | 100.0 | 83.0 | 71.5 | 2.1 | 53.9 | 14.8 | 0.8 | 4.5 | 0.5 | 4.0 | 0.0 | 0.0 |
| 15 | 100.0 | 0.2 | 0.1 | 0.0 | 0.1 | 0.0 | 0.0 | 0.1 | 0.0 | 0.0 | 0.0 | 0.0 |
| 16 | 100.0 | 0.0 | 0.0 | 0.0 | 0.0 | 0.0 | 0.0 | 0.0 | 0.0 | 0.0 | 0.0 | 0.0 |
| 17 | 100.0 | 73.6 | 55.4 | 2.4 | 46.7 | 5.9 | 0.5 | 12.8 | 0.6 | 12.1 | 0.1 | 0.0 |
| 18 | 100.0 | 7.3 | 4.2 | 0.4 | 2.1 | 1.6 | 0.1 | 1.6 | 0.9 | 0.7 | 0.0 | 0.0 |

CM - Central Memory; E - Effector (Temra); EM - Effector Memory; N - Naïve

**Table S13.** PD-L1 expression levels on T cells and relative abundances of PD-L1-positive T cells in mouse blood taken at the end of the study.

| Mouse ID | PD-L1 MFI T Cells<sup>a</sup> | PD-L1 MFI CD4<sup>+</sup> Cells<sup>a</sup> | %PD-L1<sup>+</sup> CD4<sup>+</sup> Cells<sup>b</sup> | PD-L1 MFI CD8<sup>+</sup> Cells<sup>a</sup> | %PD-L1<sup>+</sup> CD8<sup>+</sup> Cells<sup>b</sup> |
| --- | --- | --- | --- | --- | --- |
| 01 | 118 | 117 | 0,088 | 146 | 0,32 |
| 02 | 35,8 | 33,1 | 0,12 | 53,7 | 0,17 |
| 03<sup>c</sup> | 19,7 | -70,8 | 0 | -7,15 | 0 |
| 04<sup>c</sup> | -427 | N/A | 0 | -267 | 0 |
| 05 | 125 | 142 | 0,26 | 80,7 | 0,15 |
| 06 | -369 | -352 | 0,28 | -417 | 0,13 |
| 07 | 102 | 98,8 | 0,15 | 189 | 0 |
| 08 | -436 | -437 | 0,06 | -456 | 0 |
| 09 | -382 | -350 | 0 | -441 | 0 |
| 10 | 23,2 | 16,1 | 0,17 | 78,9 | 0,44 |
| 11 | -368 | -370 | 0,047 | -366 | 0,078 |
| 12<sup>c</sup> | 54,6 | N/A | 0 | -53,7 | 0 |
| 13 | 71,7 | 140 | 0 | -29,5 | 0 |
| 14 | 15,2 | 8,94 | 0,061 | 115 | 0,14 |
| 15<sup>c</sup> | -287 | -263 | 0 | -799 | 0 |
| 16<sup>c</sup> | 183 | N/A | 0 | N/A | 0 |
| 17 | 19,7 | 20,6 | 0,064 | 8,04 | 0,016 |
| 18 | 78 | 108 | 0,46 | 29,5 | 0 |

MFI - median fluorescence intensity
<sup>a</sup> Expressed as median fluorescence intensity relative to an unstained control sample.
<sup>b</sup> Abundance is relative to total CD4<sup>+</sup> or CD8<sup>+</sup> T-cell population.
<sup>c</sup> Sample contained insufficient T-cells for measuring PD-L1 expression.

## Supplementary Protocol: Isolation of Hematopoietic Stem Cells from Umbilical Cord Blood

Ensure that the working environment and solutions used in this procedure are clean and sterile. Solutions should be acclimated to room temperature (RT), unless specified otherwise in the protocol.

While the choice of density gradient medium is left up to the reader, this protocol is based on the usage of Lymphoprep™ (Cat.No. 07861, Stemcell Technologies, Canada). Lymphoprep™ should be protected from long exposure to light.

### 1 – Isolation of Mononuclear Cells (MNCs) by Density Gradient Centrifugation

1) Determine the total volume of the umbilical cord blood.
2) Aliquot a volume of density gradient medium equal to the volume of the umbilical cord blood into 50 mL conical centrifugation tubes (henceforth referred to as “50 mL tubes”). The maximum volume of density gradient medium in each 50 mL tube should not exceed 15 mL.
3) Dilute the umbilical cord blood using an equal volume of phosphate-buffered saline (PBS) or a 0.9% w/V solution of NaCl (saline).
4) Gently dispense the diluted umbilical cord blood onto the top of the density gradient medium. The volume of added blood should equal to double the volume of the density gradient medium.
5) Centrifuge the samples (400 g, 30 min, RT, with acceleration set to minimum and brakes disabled).
6) Transfer the MNCs forming the cloudy layer between the plasma and the density gradient medium into new 50 mL tubes. Ensure the maximum possible recovery of MNCs. Avoid pooling MNCs from different tubes.
7) Resuspend the MNCs in each 50 mL tube to 45 mL of total volume by adding PBS/saline and gently inverting the tubes multiple times.
8) Centrifuge the MNC suspensions (500 g, 5 min, RT).
9) Remove the supernatant by aspiration. Avoid decanting.
10) Resuspend the pelleted MNCs by adding 2 mL of PBS/saline and gently vortexing.
11) Add 20 mL of 1x Red Blood Cell Lysis Buffer (Cat.No. TNB-4300, Cytek Biosciences, USA) to each MNC suspension.
12) Gently invert the tubes multiple times to mix.
13)Incubate the MNCs in 1x Red Blood Cell Lysis Buffer (8 min, RT, protected from light).
14) Add magnetic-activated cell sorting (MACS) buffer (solution of 2 mM EDTA and 0.5% w/V bovine serum albumin in PBS, pH 7.2) into the MNC suspensions for a total suspension volume of 45 mL.
15) Gently invert the tubes multiple times to mix.
16) Centrifuge the MNC suspensions (300 g, 10 min, RT).
17) Remove the supernatant by aspiration. Avoid decanting.
*The MNCs can be frozen at this step for purification of CD34^+^ cells at a later timepoint. Once thawed, pooled and centrifuged, continue with the protocol from step 19.
18) If there are multiple tubes containing MNCs, resuspend each pellet in 1 mL of MACS buffer by pipetting and pool all of the MNC suspensions into a new 50 mL tube.
*. From this point in the protocol onwards, work with cold solutions (4°C) and keep cells in a cold environment, unless specified otherwise.
19) Resuspend the MNCs in 40 mL of MACS buffer.
20) Filter the MNC suspension through a cell strainer with a pore size of 30-40 µm into a new 50 mL tube.
21) Determine the cell count.
22) Centrifuge the MNC suspension (300 g, 10 min, 4°C).
23) Remove the supernatant by aspiration. Avoid decanting.

### 2 – Purification of CD34^+^ Human Hematopoietic Stem Cells

24) Resuspend the MNCs in MACS buffer. The total volume of the MNC suspension should be 300 µL if there are 1 x 10^8^ MNCs or fewer. For higher cell counts, scale the suspension volume proportionally so there are 1 x 10^8^ MNCs per 300 µL of suspension. During the purification of hematopoietic stem cells, scale the volumes of used reagents in the same manner. * The following CD34^+^ cell purification protocol has been adapted from that of the CD34 MicroBead Kit (human) (Cat. No. 130-046-703, Miltenyi Biotec, Germany).
25) For every 1 x 10^8^ cells, add 100 µL of human FcR blocking reagent into the MNC suspension.
26) For every 1 x 10^8^ cells, add 100 µL of CD34 microbeads into the MNC suspension.
27) Mix the MNC suspension by vortexing.
28) Incubate the MNC suspension on ice for 30 minutes, gently vortexing the suspension every 10 minutes.
29) For every 1 x 10^8^ cells, add 5 mL of MACS buffer to the MNC suspension.
30) Centrifuge the MNC suspension (300 g, 10 min, 4°C).
31) Remove as much of the supernatant as possible by aspiration. Avoid decanting.
32) For every 1 x 10^8^ cells, add 500 µL of MACS buffer to the MNC pellet.
33) Resuspend the pelleted MNCs by gently vortexing.
34) Move an aliquot of the MNC suspension containing 2 x 10^6^ cells into a separate tube for later flow cytometric analysis. Keep the aliquoted “pre-MACS” sample of MNCs cold.
35) Place an LS column into a MACS separator.
36) Rinse the LS column with 3 mL of MACS buffer and discard the flow-through.
37) Apply the MNC suspension onto the LS column and collect the flow-through fraction.
38) Wash the LS column with three 3 mL portions of MACS buffer and collect the flow-through fraction into the same tube as in step 37.
39) Move the LS column out of the magnetic field of the MACS separator and place it onto a 15 mL conical centrifugation tube.
40) Add 5 mL of MACS buffer onto the column and immediately force the added liquid through the LS column using the LS column’s plunger, collecting the eluate (the hematopoietic stem cell (HSC)-enriched fraction) into the 15 mL tube.
41) Perform steps 35-40 on the HSC-enriched fraction using a second LS column. Use the tube used in step 37 for the collection of unenriched flow-through fraction.
42) Determine the cell count in the flow-through fraction and the count and viability of the cells in the HSC-enriched fraction.
43) Move an aliquot of 1-2 x 10^6^ cells from the flow-through fraction and an aliquot of 1.5-2.5 x 10^4^ cells from the HSC-enriched fraction into separate tubes for flow cytometric analysis.
44) Keep the HSC-enriched fraction cold while determining HSC purity in cell aliquots via flow cytometry.

### 3 – Determination of the Purity of Isolated CD34^+^ Human Hematopoietic Stem Cells

45) Centrifuge the aliquots of pre-MACS, flow-through and HSC-enriched cells (450 g, 5 min, RT).
46) Remove as much supernatant as possible while minimizing cell loss.
47) Resuspend the pelleted cell aliquots. For every 5 x 10^6^ cells, rounded up, add 50 µL of PBS to the pellet.
48) Separate out an aliquot from the pre-MACS cell sample for use as an unstained control.
49) Stain the cell aliquots using an anti-human CD34 antibody (Cat.No. 130-098-140, Miltenyi Biotec, Germany).
50) Incubate antibody-stained cells for 10 minutes in a refrigerator (4°C).
51) Centrifuge the stained cell aliquots (450 g, 5 min, 4°C).
52) Aspirate supernatant.
53) Resuspend the cells in 250 µL of MACS buffer.
54) Acquire data on a flow cytometer.

### Supplementary Protocol: Preparation of Blood Samples for Spectral Flow Cytometry Analysis 1 – Thawing and Washing

1) Thaw the cryovials containing the mouse blood samples fixed using Stable-Lyse2 (Cat.No STBLYSE2-250, Smart Tube, USA) and Stable-Store2 (Cat.No STBLSTORE2-1000, Smart Tube, USA) at 4°C.
2) While the samples are thawing:

a. Label one 5 mL round-bottom tube for each sample.
b. Prepare 2 mL of 0,25 mg/mL DNAse I (Cat.No. DN25, Sigma-Aldrich, USA) in Dulbecco’s phosphate-buffered saline containing Ca^2+^ and Mg^2+^ (Cat.No. D8662, Sigma-Aldrich, USA) per sample and aliquot 1 mL of the DNAse solution into each labeled 5 mL tube. Acclimate the DNAse solution to room temperature (RT).
c. Acclimate CountBright Absolute Counting Beads (Cat.No. C36950, ThermoFisher Scientific, USA) to RT. Vortex the beads thoroughly. Into each labeled 5 mL tube containing DNAse solution, add 1 x 10^4^ counting beads for every 50 µL of blood (320 µL of fixed blood) constituting that sample.
3) One cryovial at a time:

a. Pipet the contents of the cryovial gently and thoroughly to resuspend the cells.
b. Transfer the contents of the cryovial into the corresponding 5 mL tube containing DNAse solution and counting beads.
c. Add 1 mL of DNAse solution into the cryovial.
d. Wash out the cryovial with the DNAse solution and transfer the washout to the corresponding 5 mL tube.
e. Pipet gently and thoroughly to mix the contents of the 5 mL tube.
4) Incubate cells in DNAse solution for a minimum of 10 minutes.
5) Centrifuge the samples (800 g, 5 min, RT).
6) Remove supernatant by pipetting.
7) Add 1 mL of phosphate-buffered saline (PBS) to each sample.
8) Resuspend the cells by vortexing.
9) One sample at a time:

a. Pipet the resuspended cells through the filter of a correspondingly labeled filter-capped 5 mL tube (Cat.No. 352235, Corning, USA).
b. Add 1 mL of PBS to the original 5 mL tube.
c. Wash the walls of the tube by vortexing.
d. Transfer the washout through the filter of the corresponding 5 mL tube.
10) Centrifuge the samples (800 g, 5 min, RT).
11) Remove as much supernatant as possible by pipetting.

### 2 – Staining

12) Prepare FcR blocking buffer:

a. 38 µL of base buffer (2% V/V fetal bovine serum in PBS) per sample.
b. 10 µL of human FcR blocking agent (Cat.No. 130-059-901, Miltenyi Biotec, Germany) per sample.
c. 2 µL of anti-mouse-CD16/CD32 monoclonal antibody (Cat.No. 16-0161-82, ThermoFisher Scientific, USA) per sample.
13) Add 50 µL of FcR blocking buffer to each cell pellet.
14) Resuspend the cells by gently vortexing.
15) Incubate cells in FcR blocking buffer for 10 minutes at RT.
16) Add 50 µL of antibody mix to each sample.
17) Mix by gently vortexing.
18) Incubate cells in the antibody mix for 30 minutes in a fridge (4°C).
19) Add 2 mL of base buffer to each sample.
20) Mix by vortexing.
21) Centrifuge the samples (800 g, 5 min, RT).
22) Remove supernatant by pipetting.
23) Repeat steps 19-22 to perform a second cell wash.
24) Resuspend the cells by gently vortexing.
25) One sample at a time:

a. Measure the volume of the cell suspension using a pipette.
b. Adjust the volume of the cell suspension to 200 µL using base buffer.
26) Acquire the samples on an ID7000 Spectral Cell Analyzer (LE-ID7000C, Sony Biotechnology, USA).

### Supplementary Protocol: Immunohistochemical Staining of Primary Mouse Tumor Sections

#### Day 1

1) Bake the slides with the primary tumor sections at 60°C for at least 1h.
2) Cool the slides to room temperature (RT).
3) Deparaffinize and rehydrate the sections by incubation in xylene and a graded ethanol series:

a. Xylene – pass 1 (10 min, RT).
b. Xylene – pass 2 (3 min, RT).
c. Ethanol (100%) – Pass 1 (3 min, RT).
d. Ethanol (100%) – Pass 2 (3 min, RT).
e. Ethanol (96%) – Pass 1 (3 min, RT).
f. Ethanol (96%) – Pass 2 (3 min, RT).
g. Ethanol (80%) (3 min, RT).
h. Deionized water (diH_2_O) (5 min, RT).
4) Move the slides to a container of 1x Dako Target Retrieval Solution, pH 9 (Cat.No. S2367, Agilent, USA).
5) Perform heat-induced epitope retrieval. For example, by a 20-minute incubation in a microwave (Model JT366/WH, Whirlpool, USA) on a no-boil (“6^th^ sense”) setting.
6) Cool the slides in antigen retrieval solution on a benchtop for 10 minutes.
7) Further cool the slides to RT by placing the container of slides in antigen retrieval solution into a sink and pouring room-temperature diH_2_O into the container.
8) Dry the back of the slides and the area surrounding the tumor section using tissue paper.
9) Outline the tumor sections tightly using a PAP marker (Cat.No. Z627548, Sigma-Aldrich, USA).
10) Place the slides into hydration chambers containing moistened tissue paper.
11) To keep tumor sections hydrated while drying other slides, temporarily add diH_2_O onto tumor sections on dried slides.
12) Remove excess diH_2_O from the tumor sections by shaking the slides.
13) Add Dako Real™ Peroxidase Blocking Solution (Cat.No. S2023, Agilent, USA) onto the tumor sections.
14) Incubate for 10 minutes at RT.
15) Wash off the peroxidase blocking solution into a waste container by applying 1x Dako Wash Buffer (Cat.No. S3006, Agilent, USA) using a squeeze bottle. Temporarily leave a small amount of wash buffer on the tumor sections to keep them hydrated while washing other slides.
16) Remove excess wash buffer from the tumor sections by shaking the slides.
17) Add wash buffer onto the tumor sections.
18) Incubate for 5 minutes at RT.
19) Remove the wash buffer from the slides into a waste container.
20) For a second time, add wash buffer onto the tumor sections.
21) Incubate for 5 minutes at RT.
22) Wash off the wash buffer into a waste container by applying diH_2_O using a squeeze bottle. Sufficient wash buffer has been removed once the surface tension of the liquid on the slide allows the PAP marker outline to become clearly visible.
23) To keep tumor sections hydrated while washing other slides, temporarily add diH_2_O onto washed tumor sections.
24) Remove excess diH_2_O from the tumor sections by shaking the slides.
25) Add blocking solution (3% w/V bovine serum albumin (BSA) in phosphate-buffered saline (PBS)) onto the tumor sections.
26) Incubate for 45-60 minutes at room temperature.
27) Wash off the blocking solution into a waste container by applying wash buffer using a squeeze bottle. Temporarily leave a small amount of wash buffer on the tumor sections to keep them hydrated while washing other slides.
28) Remove excess wash buffer from the tumor sections by shaking the slides.
29) Add wash buffer onto the tumor sections.
30) Incubate for 5 minutes at RT.
31) Remove the wash buffer from the tumor sections into a waste container.
32) For a second time, add wash buffer onto the tumor sections.
33) Incubate for 5 minutes at RT.
34) Wash off the wash buffer into a waste container by applying diH_2_O using a squeeze bottle. Sufficient wash buffer has been removed once the surface tension of the liquid on the slide allows the PAP marker outline to become clearly visible.
35) To keep tumor sections hydrated while washing other slides, temporarily add diH_2_O onto washed tumor sections.
36) One slide at a time, remove as much excess diH_2_O as possible, avoiding damaging the tumor sections, then apply the appropriate primary antibody (diluted in 0.5% w/V BSA) to the tumor sections.
37) Incubate tumor sections in primary antibody overnight in a fridge (4°C).

#### Day 2

38) Retrieve the primary-antibody-stained tumor sections from the fridge.
39) Wash off the primary antibody solutions by tilting the slide over a waste container and applying wash buffer using a squeeze bottle.
40) Immediately place washed slides into a container of wash buffer.
41) Incubate on an orbital shaker for 5 minutes at RT.
42) Move the slides into a fresh container of wash buffer.
43) Incubate on an orbital shaker for 5 minutes at RT.
44) Wash off the wash buffer into a waste container by applying diH_2_O using a squeeze bottle. Sufficient wash buffer has been removed once the surface tension of the liquid on the slide allows the PAP marker outline to become clearly visible.
45) To keep tumor sections hydrated while washing other slides, temporarily add diH_2_O onto washed tumor sections.
46) Remove excess diH_2_O from the tumor sections by shaking the slides.
47) Apply the appropriate horseradish-peroxidase-conjugated secondary antibody (e.g. Dako EnVision+ System-HRP Labelled Polymer Anti-Mouse (Cat.No. K4001, Agilent, USA) or Dako EnVision+ System-HRP Labelled Polymer Anti-Rabbit (Cat.No. K4003, Agilent, USA)) to the tumor sections.
48) Incubate the tumor sections in secondary antibody for 30 minutes at RT.
49) Wash off the secondary antibody solutions by tilting the slide over a waste container and applying wash buffer using a squeeze bottle.
50) Immediately place washed slides into a container of wash buffer.
51) Incubate on an orbital shaker for 5 minutes at RT.
52) Move the slides into a fresh container of wash buffer.
53) Incubate on an orbital shaker for 5 minutes at RT.
54) Wash off the wash buffer into a waste container by applying diH_2_O using a squeeze bottle. Sufficient wash buffer has been removed once the surface tension of the liquid on the slide allows the PAP marker outline to become clearly visible.
55) To keep tumor sections hydrated while washing other slides, temporarily add diH_2_O onto washed tumor sections.
56) Remove excess diH_2_O from the tumor sections by shaking the slides.
57) Apply a solution of diaminobenzidine (Liquid DAB+, 2-component system (Cat.No. K3468, Agilent, USA)) to the tumor sections. Exercise caution while handling and disposing of diaminobenzidine due to its toxicity.
58) Incubate tumor sections in diaminobenzidine solution in the dark for 8 minutes at RT.
59) Thoroughly wash off the diaminobenzidine solution by tilting the slide over a toxic waste container and applying diH_2_O using a squeeze bottle.
60) To keep tumor sections hydrated while washing other slides, temporarily add diH_2_O onto washed tumor sections.
61) Remove excess diH_2_O from the tumor sections by shaking the slides.
62) Place the slides into a container of hematoxylin (Cat.No. S3301, Agilent, USA).
63) Incubate the tumor sections in hematoxylin for 10 minutes at RT.
64) Remove the slides from the hematoxylin and shake off the excess.
65) Place the slides into an empty container and wash the remaining hematoxylin off using several portions of warm tap water.
66) Place the slides into a container of diH_2_O.
67) Dehydrate the tumor sections and prepare them for mounting by dipping them 10 times into each container of a graded ethanol series and xylene:

a. Ethanol (80%)
b. Ethanol (96%) – Pass 1
c. Ethanol (96%) – Pass 2
d. Ethanol (100%) – Pass 1
e. Ethanol (100%) – Pass 2
f. Xylene – pass 1
g. Xylene – pass 2
68) Mount the slides.

